# Ligand-induced unfolding mechanism of an RNA G-quadruplex

**DOI:** 10.1101/2021.10.26.465985

**Authors:** Susanta Haldar, Yashu Zhang, Ying Xia, Barira Islam, Sisi Liu, Francesco L Gervasio, Adrian J. Mulholland, Zoë A. E. Waller, Dengguo Wei, Shozeb Haider

## Abstract

The cationic porphyrin, TMPyP4, is a well-established DNA G-quadruplex (G4) binding ligand that can stabilize different topologies via multiple binding modes. However, TMPyP4 has completely opposite destabilizing and unwinding effect on RNA G4 structures. The structural mechanisms that mediate RNA G4 unfolding remains unknown. Here, we report on the TMPyP4-induced RNA G4 unfolding mechanism studied by well-tempered metadynamics (WT-MetaD) with supporting biophysical experiments. The simulations predict a two-state mechanism of TMPyP4 interaction via a groove-bound and a top-face bound conformation. The dynamics of TMPyP4 stacking on the top tetrad disrupts Hoogsteen H-bonds between guanine bases resulting in the consecutive TMPyP4 intercalation from top-to-bottom G-tetrads. The results reveal a striking correlation between computational and experimental approaches and validate WT-MetaD simulations as a powerful tool for studying RNA G4-ligand interactions.

## Introduction

Guanine rich sequences in single-stranded DNA/RNA can self-associate, in a co-planar, cyclic arrangement called G-tetrads.^1, 2^ Each tetrad comprise of four guanine bases, stabilized by eight Hoosteen hydrogen bonds reinforced by π and σ bonds.^3, 4^ The G-tetrads can stack on top of one another, held together by π-π stacked non-bonded attractive interactions to form a G-quadruplex (G4).^5, 6^ G4 structure is further stabilized by the presence of mono or divalent cations as they coordinate between two successive G-tetrads and shield the electrostatic repulsion between the carbonyl oxygens of guanines.^5^ G4-DNA structures are polymorphic and can adopt varied topologies influenced by strand stoichiometry (one to four), polarity (parallel, antiparallel, hybrid), glycosidic conformation (syn/anti), intervening length of loops and guanine stretches.^7^ G4-RNA structures, however, are limited to predominantly parallel topology, primarily due to the anti-conformation of the glycosidic bonds and the presence of an additional 2’-OH group that contributes to enhanced hydrogen bond networks.^8, 9^

G4s are widespread throughout the human genome. Extensive human genomic analyses have identified >500,000 DNA and >6000 RNA putative G-quadruplex forming sequences (PQFS).^10–13^ They are found to colocalize with functional regions of the genome including specific sites such as telomeres and promoter regions of genes.^14–18^ Their presence has important implications in regions involved in telomere maintenance and persistent DNA damage response in ageing.^19^ Human telomeric DNA d(TTAGGG) and the non-coding TERRA RNA r(UUAGGG) can both adopt characteristic G4 structures and are important participants in telomere biology and epigenetic regulation.^9, 20^ Exhaustive mapping of the genome-wide location of DNA replication origins have revealed that ∼90% of start sites contain the PQFS.^21^ Such high genomic distribution of PQFS in gene transcription start sites is indicative of G4-mediated role in regulating the process of transcription.^22, 23^ Furthermore, significant enrichment of PQFS in the vicinity of somatic copy-number alterations breakpoints is an epigenetic determinant driving tissue-specific mutational landscapes in cancer.^24^ New functional roles of G4-RNA have also been reported in the regulation of RNA expression in mitochondria,^25^ in phase separation mechanisms leading to the formation of membrane-less organelles,^26, 27^ and epitranscriptomics.^28^ A high frequency of PQFS has also been reported in region specifying the 5’-UTR of the encoded mRNAs, suggesting G4-mediated role in regulating translation.^29, 30^ Moreover, the formation of G4-DNA in cells has been shown to be linked to replication and transcription, and that of G4-RNA with translation.^31^ Based on these observations, it has thus been proposed that G4 (un)folding ***in vivo*** could possibly represent another layer of non-genetic, but structural regulation of gene expression.^32^ Besides the human genome, PQFS are also present extensively in bacteria,^33^ and viruses.^34^

The involvement of G4s in several biological processes has made them a potential target for therapeutic intervention. The structural features of G4s have been exploited to increase their thermodynamic stability via induced formation by small molecules.^15, 35^ G4s with high thermodynamic stability are obstruction to processivity by the cellular replicative machinery. This was the central idea behind the design of G4s stabilizing agents targeting telomere maintenance,^36^ or associated suppression of transcriptional activation in proliferative cancer cells.^37^ Most of the G4 interacting small molecules present in the literature focus on the stabilization of G4 structure.^38–40^ On the other hand, the importance of destabilizing G4 structures has been underexplored. Destabilization of G4 structures have been shown to enhance translational efficiency in the FMR1 gene and the 5’ UTR of the FMR1 mRNA in Fragile X syndrome.^41, 42^ Furthermore, a genetic loss or age-related changes in G4 modulating proteins, compounded by over representation of G4s have been shown to accelerate brain aging and foster neurological disorders.^43^ Thus, destabilization of G4 structures can be another means of controlling gene expression or find applications in treating age-associated neurobiological disorders. However, it has not been trivial to reliably assess G4 destabilization by small molecules due to the lack of standard assays and protocols.^32^ It is only recently (while this manuscript was under review) that Monchaud and co-workers have published a G4-helicase based destabilization assay.^44^ Nevertheless, there are small molecules that have been reported to destabilize G4s. For example, TMPyP4 disrupted the G4 structure in the Fragile X FMR1 gene,^42^ and in the MT3 endopeptidase mRNA sequence;^45^ a triarylpyridine derivative disrupted G4 in the cKit-1 and 2 sequences;^46^ an anthrathiophenedione,^47^ and a stiff-stilbene derivative was shown to unfold a sodium form of telomeric G4.^48^

The omnipresence of G4 structures in a cell-based setting presents a formidable challenge to stabilize or unwind a specific topology. The formation and dissolution of G4s have been studied by a wide variety of biophysical and chemical probe methods.^45, 48–50^ Several G4-ligand complexes have been characterized by crystallographic and NMR studies.^51–57^ Recently, the DEAH/RHA helicase DHX36-G4 complex structure laid the structural foundation to explore the G4 unfolding mechanism by a helicase.^58^ Besides, numerous helicases that unwind G4 have been identified and are being used as molecular tools to study G4 unwinding.^32^

In the absence of crystal structures, computational methods have been an indispensable tool to study such processes. Recently, Moraca et al. showed the binding mechanism of berberine (a polycyclic aromatic compound) to human telomeric G4-DNA and its stabilization through a combined effort using well-tempered metadynamics (WT-MetaD) simulation and steady-state fluorescence spectroscopy experiments.^59^ In another study, O’Hagan et al. studied the reversible unfolding mechanism of G4-DNA by a photo responsive stiff-stilbene ligand in the presence of sodium buffer using WT-MetaD simulations, circular dichroism and NMR spectroscopy.^48^ In particular, this study used human telomeric antiparallel G4-DNA to investigate unfolding. One common feature that was identified while looking at the ligand-complex structures that the binders are mostly one or more polycyclic and planar aromatic chromophores, able to engage in π-π stacking interactions with the terminal G-tetrads, and a positive charge that is necessary to interact with the DNA backbone phosphate groups.^40^ Specifically, this study also showed that the unfolding of G4-DNA initiated from the groove via the formation of the π-π stacking interaction between the stilbene moiety and G-tetrads.^48^

Among all classical G4 interacting ligands, TMPyP4 is a paradox that exhibits both G4 stabilizing and destabilizing properties. TMPyP4 has been shown to stabilize G4 structures and exert anti-tumor,^60, 61^ and anti-viral activity.^62^ In the NMR structure (PDB id 2A5R), TMPyP4 was reported to stabilize the c-Myc Pu24I by stacking with the surface of 5’-quartet.^57^ Neidle and co-workers explored the interactions between TMPyP4 and biomolecular human telomeric quadruplex (PDB id 2HRI), in which, TMPyP4 exhibits an alternative binding mode by stacking with the loop TTA nucleotides instead of the G-quartet.^63^ However, TMPyP4 is also reported to have a completely opposing effect on G4-RNA structure and unwinds them.^45, 49^ Until now, there is no structural insight on how TMPyP4 or any small molecule unwinds G4-RNA structures.

We approached this issue by studying the interactions between TMPyP4 and an RNA G4 forming sequence (PQS18-1; r(GGCUCGGCGGCGGA)) from the non-coding region of Pseudorabies virus (PRV).^64^ The structure of PQS18-1 (PDB id 6JJH, 6JJI) has been published recently.^65^ It is a bimolecular, all-parallel stranded G4-RNA consisting of 4-stacked tetrads with three K+ ions positioned along the central axis. The structure contains two molecules of TMPyP4 ligand bound to the G4; one intercalated at the 3’ end between the top G-tetrad plane and an A-diad formed from A14:A14’ bases and one external to the G4-RNA in a stacked arrangement between a pair of uracils (U4:U4’) and a pair of cytosines (C11:C11’) (Figure 1).

**Figure 1:**
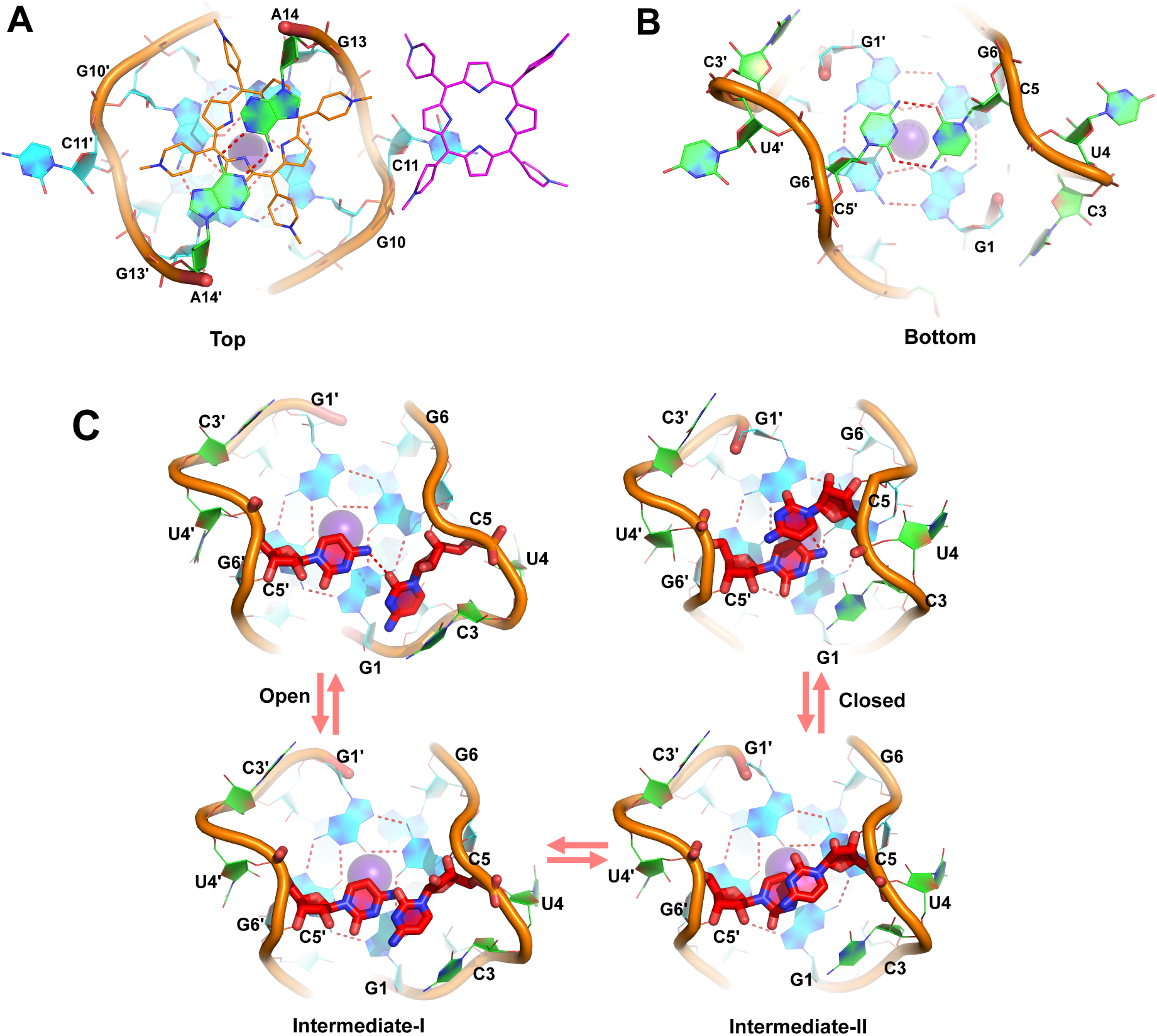
**Crystal structure of the RNA-TMPyP4 complex (PDB id-6JJH).** (a) the top-face view highlighting the two different bound states of TMPyP4. TMPyP4 shown in orange is sandwiched between A14, A14’ and guanine bases, G10, G10’, G13, G13’, from the first G-tetrad. TMPyP4 shown in magenta is π-π stacked with C11 base. (b) the bottom-face view of the RNA-TMPyP4 complex highlighting a H-bond network between the capped bases C5 and C5’ in an open like conformational arrangement. (c) The dynamic equilibrium between open and closed conformational transitions via two intermediate states (I and II). Both the C5 and C5’ bases are highlighted for better visuals with default colors of the atoms (Carbon in red, Oxygen in light red and Nitrogen in blue).

Herein we report a computational study of the unfolding of an RNA G4 topology by porphyrin TMPyP4 ligand from a well-balanced WT-MetaD enhanced sampling simulation. Our simulation data (∼1.5 μs) showed that TMPyP4 binds on the top-face as well as in the groove side of the RNA G4 mimicking the X-ray crystal structure binding poses.^65^ Moreover, extending the simulation to a further 1.1 μs (a total of ∼2.6 μs) highlighted the complete unfolding of the RNA G4 topology. Further to validate our theoretical results and assess the reliability of the method used, we performed several experiments. Through WT-MetaD simulations, circular dichroism (CD), ultraviolet-visible (UV-vis) absorbance, fluorescence titrations (FRET), and Isothermal Titration Calorimetry (ITC) experiments, we hypothesize that the unfolding process could be divided into three states: 1) an initial binding between the ligands and RNA G4, 2) dynamic movement of the intermediates or formation of the intermediate complex through opening of the G-tetrads via intercalation and 3) unfolding. Finally, our results indicate that computational simulations can predict ligand-induced unfolding of G4s with precision. The WT-MetaD data complements experimental findings and provides extensive structural information for the TMPyP4 mediated RNA G4 unfolding process, therefore it can be used as a useful tool for investigating G4/ligand interactions.

## Results

### Unbiased MD simulations

To gain insights into TMPyP4 ligand binding to RNA G4, we carried out unbiased classical MD simulations of both the native RNA G4 and all available TMPyP4-PRV PQS18-1 RNA G4 complexes (PDB id 6JJH, 6JJI).^65^ Crystal structures revealed a top-face bound state of TMPyP4 where it is sandwiched in between the top fraying base A14 and A14’ and the bases from the first G-tetrad such as G10, G10’, G13 and G13’.^65^ In the second bound state, TMPyP4 is mainly solvent exposed and stacked on the C11 base of RNA G4 loop (Figure 1a). We performed 1 μs (x3) of classical MD simulation of both the native and TMPyP4 groove bound states to test the stability of these systems. Further, we carried out Principal Component Analysis (PCA) of all the systems investigated and then compared the results to understand the dynamic behaviour of the RNA G4 structure. PCA analysis on the native G4 shows that the RNA G4 is highly flexible in water and can rapidly change conformations between open and closed states. At the start, both the C5 and C5’ bases are positioned adjacent to one another on the top plane of the terminal G-tetrad, (Figure 1b, 1c) resulting in the backbones of chain A and chain B to spread out. We consider this as the open like conformation as found in the crystal structure (Figure 1b, Bottom). In the closed state, both the C5 and C5’ bases are found to be stacked on top of each other, and the RNA G4 shrinks by bringing the backbone of chains A and B close to each other relative to the open state. Intermediate states (I and II, Figure 1) shows how the C5 and C5’ bases transform their position from open-to-closed states (Figure 1c).

Further, MD simulation demonstrates that the top-face bound state is stabilized in the open like conformation whereas the groove-bound state resembles the closed conformation. Figure 1 shows the sequence of transformation from open-to-closed conformation and vice-versa. (See Supplementary Information for more details on the discussion of PCA analysis of native RNA G4 and its complexes).

### Free energy calculations using well-tempered metadynamics simulations

To study the complete binding and unfolding of TMPyP4 ligand to RNA G4, we performed WT-MetaD simulation. Metadynamics (MetaD) is an enhanced sampling simulation technique which allows simulating long time scale events considered as rare events such as protein-ligand binding,^66^ protein folding/unfolding mechanism,^67^ in a reasonable computational time cost.^68^ At the end of the simulation, the free energy landscape of the simulated process of interest can be computed using the history-dependent biasing potential which was added during the simulation on a chosen degree of freedom called Collective Variables (CVs).^68^ This technique has already been successfully used by us and also by few other research groups to simulate biological processes like folding/unfolding mechanism of RNA tetraloops,^69, 70^ G4-ligand binding^48, 71^ and also in the materials science such as binding of small ligands to the surfaces. ^72, 73^

In the present study, MetaD simulation was used to (a) explore the available binding modes of TMPyP4 ligand around the RNA G4, (b) identify a possible unfolding mechanism of the RNA G4 and (c) compare the observed aforementioned phenomena to the available experimental data. Two collective variables were used for WT-MetaD to study binding and unbinding of TMPyP4 ligand around the RNA G4. These are the distance (D) between the center of mass (COM) of the ligand to the COM of the middle G-tetrad in the RNA G4 and a torsion (T) angle between the ligand and the G4. The torsion angle between the ligand and G4 was measured as the two points from ligand to two points from the RNA G4 (see Table S1 in Supplementary Information for details). The simulation took over 2.6 μs to converge with the following protocol: first, a single WT-MetaD simulation is performed to reach a semiquantitative convergence where we have sampled all the possible free energy minima for the ligand binding to RNA G4. Particularly, two separate binding conformations (basins) are sampled and they are the top-face and groove-binding conformations. To further accelerate the sampling of these conformations, four consecutive walkers are placed along the path from successive basins using the multiple walkers technique for rigorous sampling (see below). A two-dimensional representation of the free energy surface (FES), as a function of D and T, is shown in Figure 2. Further, to have a quantitatively well-characterized free energy profile, various recrossing events between the different states such as bound and unbound states, visited by the system should be seen. To provide a picture of the convergence of the binding free energy estimation, the free energy difference between the bound and unbound states was computed as a function of the simulation time (Figure S1). The estimate of free energy of binding of TMPyP4 to RNA G4 converges to -10.5 kcal/mol (ΔG_cal_), which is close to the experimentally obtained binding free energy (ΔG_expt_) of -10.2 kcal/mol (Table 1).

**Figure 2:**
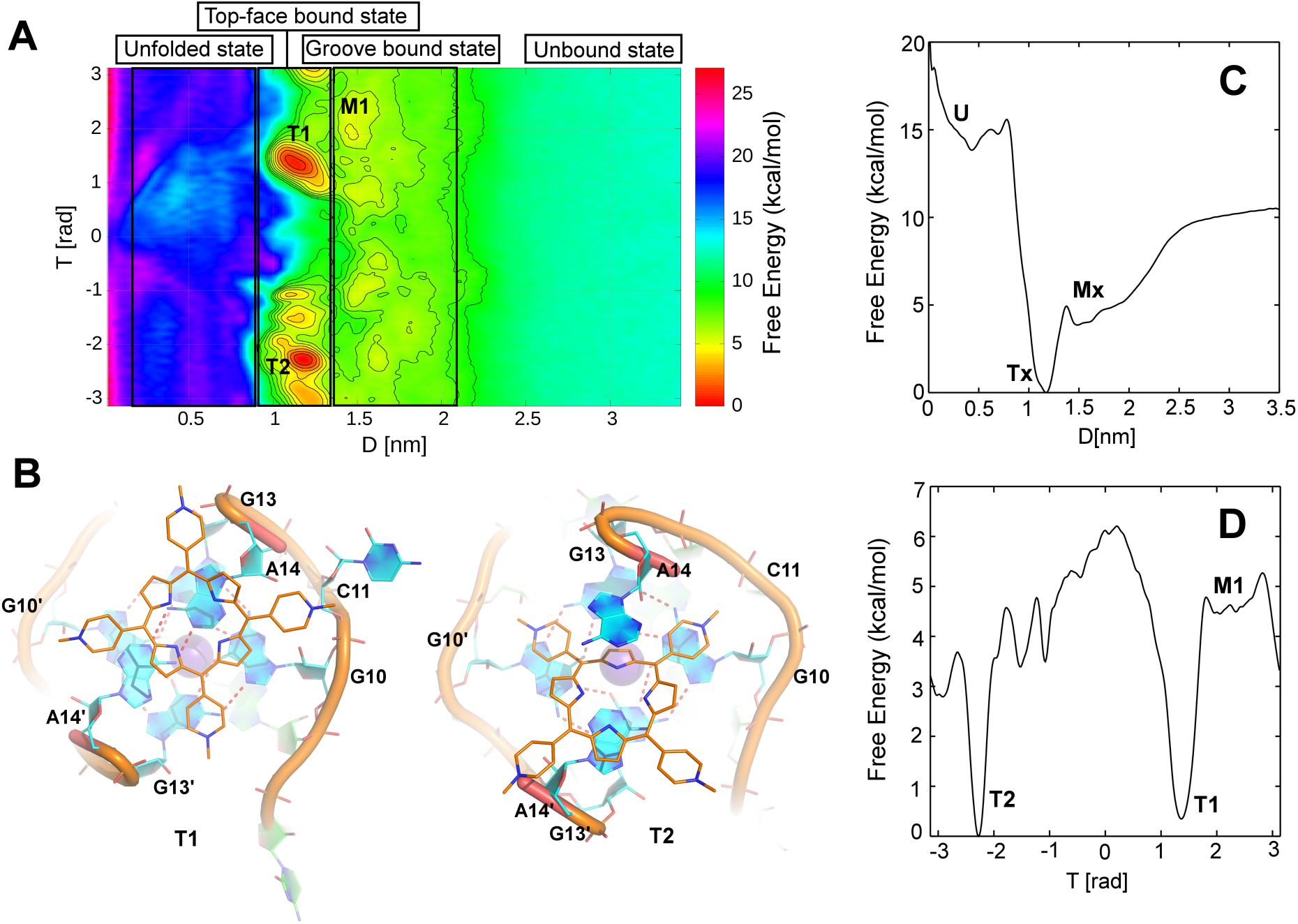
**Free energy calculations of TMPyP4 binding to PQS18-1 RNA G4.** (a) Free energy surface (FES) of binding of TMPyP4 to RNA G4 topology illustrated as a function of distance (D) and torsion (T) collective variables. Each box in the FES represents a different state of RNA such as TMPyP4 bound to the RNA top-face (0.9 ≤ D ≤ 1.4), TMPyP4 bound to the RNA Groove (1.4 ≤ D ≤ 2.1), Unbound state, (>3.0) and TMPyP4 mediated RNA unfolded state (U) (0.0 ≤ D ≤ 0.9). Two most pronounced basins are found in the top-face bound state, T1 and T2. (b) The most populated clusters are shown for both the top-face bound states. In T1, TMPyP4 binds on the extreme top of the RNA interacting with A14 and A14’ fraying bases whereas, in T2, TMPyP4 is found to be slightly tilted relative to the T1 pose, interacting with first G-quartet bases, G10, G10’ G13, and A14 and A14’ nucleotide bases and mimicking the native crystal TMPyP4-RNA G4 bound complex. (c) The one-dimensional potential of mean force (PMF) is plotted as a function of distance (D) in nm, showing the ΔG_cal_ = -10.5 kcal/mol. Free energy basins such as Tx, Mx, and U represent the corresponding top-face, groove-bound and unfolded states. (d) The one-dimensional PMF plotted as a function of torsion (T) CV in radians, illustrating the absolute free energy difference between T1 and T2 basin.

**Table 1:** Thermodynamic parameters of ITC experiments

| Experiment | Ka (M<sup>-1</sup>) | $\Delta H$ (cal/mol) | $\Delta S$ (cal/mol/deg) | n | $\Delta G_{\text{expt}}^*$ (kcal/mol) | $\Delta G_{\text{calc}}$ (kcal/mol) |
| --- | --- | --- | --- | --- | --- | --- |
| RNA<br>PQS18-1<br>with<br>TMPyP4 | 5.74(±0.46) | -2.17(±0.09) | -46.5 | 1.24±0.02 | 7.86 | 6.7 |
| | 2.83(±2.67)<br>$\times 10^7$ | -6.87(±4.08)<br>$\times 10^2$ | 31.8 | 0.32±0.04 | 10.17 | 10.5 |

The $\Delta G_{\text{expt}}$ is calculated from the equation of $-RT\ln K_a$ where R, T and $K_a$ are the universal gas constant, standard temperature of the reaction and binding constant, respectively.

### TMPyP4-RNA G4 interactions: Top-face and major groove binding modes

The available crystal structure confirmed the top-face and groove-binding modes of TMPyP4 to RNA G4.^65^ The two-dimensional FES (Figure 2a) can distinguish different states including top-face bound state, groove-bound state, unbound state and the unfolded state. Top-face bound states are represented as Tx where x corresponds to the number of bound states based on their position and orientation (Figure 2). The top-face bound states are assigned within the distance 0.9 ≤ D ≤ 1.4 nm in the D versus T curve. In basin T1, a cluster analysis using an RMSD cutoff value of 0.17 nm indicated that TMPyP4 acquire a top-face binding conformation (with 93.4 % population) where the major contribution comes from the stacking interaction with both the top A14 and A14’ nucleotide bases (Figure 2). In particular, all four pyrrole rings are involved in the π-π stacking interaction with the aforementioned bases (see T1, Figure 2). We note that few ionic interactions are also seen between the pyridinium cation and negatively charged oxygen atoms of the phosphate backbone of RNA G4. Basin T2 shows a rather tilted top-face binding conformation where TMPyP4 is intercalated in between the A14 and A14’ fraying bases. Further, TMPyP4 is also seen to partially stack with G10, G13 and G10’ bases with a population of 75.6 % as calculated from the cluster analysis. This pose is found to be very similar to the available top-face TMPyP4-RNA G4 binding pose in the X-ray crystal structure (Figure 1). Energetically, both the top-face bound states (T1 and T2) are found to be very similar in their binding free energy with a difference being 0.3 kcal/mol (Figure 1). Although the present WT-MetaD simulation is not able to capture the exact crystal structure top-face binding conformation but has been quite successful to sample the near-native binding conformations (Figure 1). Further, Principal Component Analysis (PCA) on the top-face binding pose revealed that TMPyP4 binding on the top-face allows the RNA to adopt an open like conformation (Figure S3). Moreover, the PCA analysis is also performed on the native RNA itself to differentiate the open to closed conformational changes and compare it with the bound states (Figure S2).

Major groove binding modes are represented as Mx where x accounts for the number of TMPyP4 groove bound states differing in their position interacting with different nucleotide bases around the RNA G4. The groove bound states are assigned within the distance 1.4 ≤ D ≤ 2.1 nm in the D versus T curve. A cluster analysis on the most stable basin (M1, Figure 1 and Figure S5) in the groove binding region revealed a high degree of heterogeneity on the ligand binding position i.e. towards different binding sites; since the RNA G4 has a top-face and four groove binding sites. Three equally populated clusters from the basin M1 were obtained (Figure S5). TMPyP4 is either bound to the top-face of the RNA G4 interacting with fraying adenine (A14 and A14’) bases or in the groove site binding individually to C11 and C11’ bases (see Supporting Information for more details). Thus, the groove-binding sites have not well been separated by the FES portrayed with D and T CVs in Figure 2.

To address this issue, we explored several variants of CVs. Following a previous study by O’Hagan et al.,^74^ we have adopted two new CVs such as ligand position in the x- and y-axis (taken as the vector of the distance in the x and y direction i.e. d.x and d.y, respectively). This condition does only apply when the RNA principal axis is aligned with the z-axis (Figure S6). The vectors are able to describe the possible groove-binding modes of TMPyP4 since the groove sides lie in the x and y plane (Figure 3). The major advantage of using these two CVs is to separate each groove bound-state in the four-fold symmetry. These CVs are also able to separate the top-face bound state as well. Looking at the FES, four additional basins are found along with T, the most stable one which represents the top-face TMPyP4 bound state (Figure 3). In basin M1, a cluster analysis revealed that TMPyP4 adopts a groove binding conformation in which it is mostly interacting with C11 base via π-π stacking interaction. This pose agrees well with our crystal structure (PDB id 6JJH).^65^ In particular, one of the pyrrole rings from TMPyP4 is involved in the stacking interaction. We carried out a 1 μs (x3) unbiased MD simulation with ligand-bound in this pose. The TMPyP4 remained stacked with C11 and no significant deviation from starting structure was observed indicating that this pose was stable (Figure S7). Further, a PCA analysis on the MD data of RNA G4 with TMPyP4 in the groove-bound state suggests that the RNA G4 topology remains in a closed conformation for the M1 binding mode (Figure S4). This finding has great importance on the TMPyP4-mediated unfolding mechanism of the RNA G4 topology.

**Figure 3:**
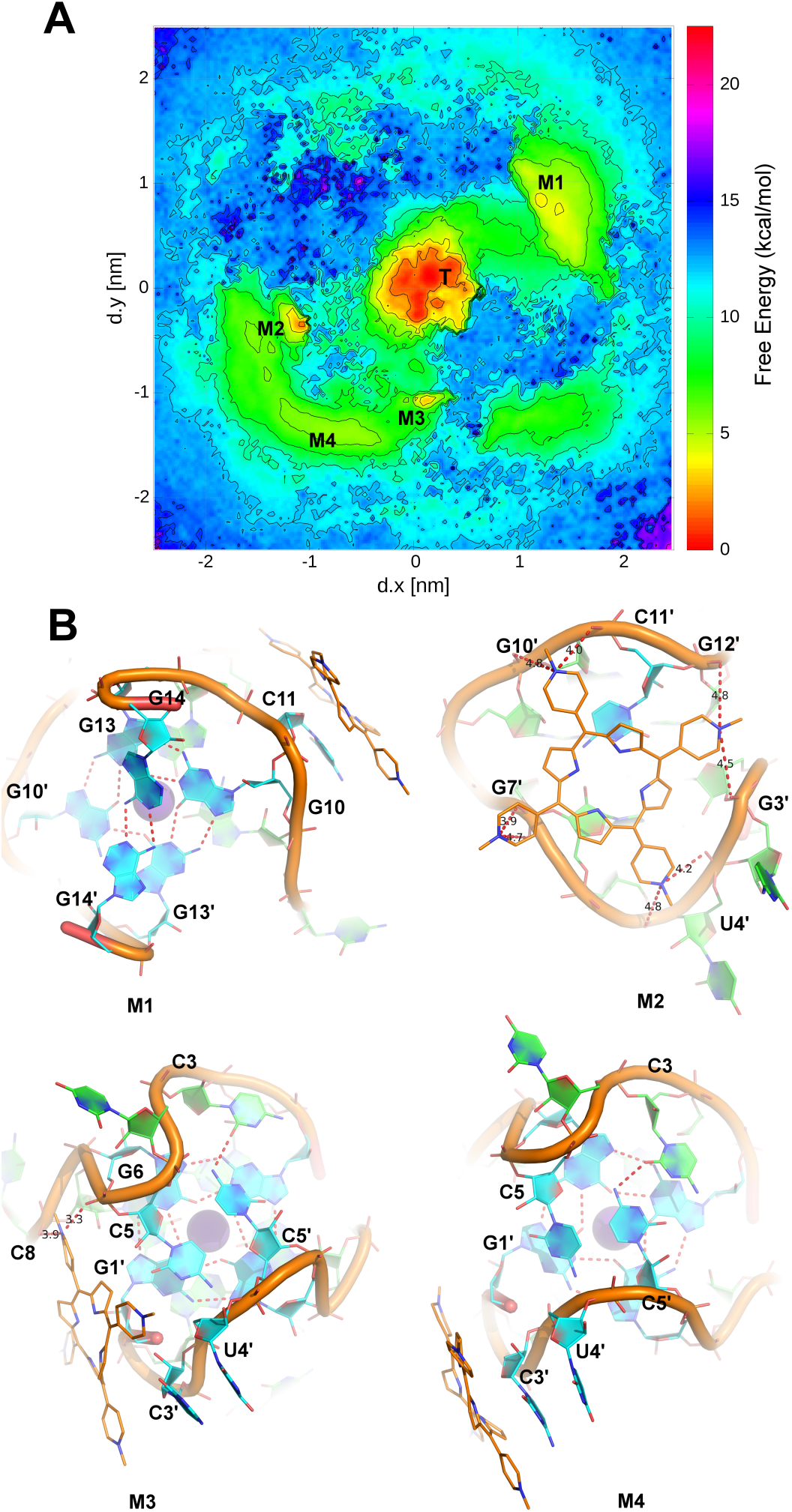
**Free energy surface of groove binding states.** (a) Free energy surface plot considering d.x and d.y as the two major collective variables (CVs). The CVs were able to capture the TMPyP4-RNA G4 groove bound state. Basin T at the centre represents the top-face binding conformation whereas Mx corresponds to the groove binding conformations. (b) Basin M1 and M2 represent the X-ray crystal structure-like groove bound states where TMPyP4 interacts with C11 and C11’ bases, respectively. In basin M3, TMPyP4 partially interacts with C3’ base, in addition to the negatively charged oxygen atoms from the RNA backbone with its positively charged pyridine nitrogen’s. Basin M4 is the least stable free energy basin where TMPyP4 mainly interacts with C3’ base.

A cluster analysis was performed for basin M2 where TMPyP4 remains on the opposite side to C11-C11’ base stacking interactions. In particular, several electrostatic interactions are established between pyridinium cation and backbone oxygen atoms of the G3’, G12’, G10’, G7’ and U4’ nucleotides. The stability of this pose is again validated with ∼1 μs (x3) unbiased MD simulation. As no significant deviation from the starting structure was observed, we concluded that it is a stable binding pose (Figure S7). This pose was similar to the available X-ray crystal structure ^65^ since C11’ base is equivalent to C11 due to the observed symmetry between both RNA strands (chain A and chain B). Basin M3 is a groove-binding pose involving mainly electrostatic interactions and partially one of the ligands pyridine ring interacting with C3’ base via π-π stacking interactions. Moreover, in this conformation, one of the pyrrole rings of TMPyP4 makes stacking interaction with G1’ sugar pucker ring. Furthermore, an electrostatic interaction is established between the pyridinium cation and RNA backbone oxygen atoms of C8 and G6 nucleotides. In basin M4, a solvent-exposed conformation of TMPyP4 is observed where it interacts with the C3’ base via π-π stacking interaction. In particular, one of the pyrrole ring from the porphyrin moiety and the pyridine ring is involved in the π-π stacking interaction with C3’ base. The only difference between basin M3 and M4 is that in the present conformation, TMPyP4 has slightly shifted towards C3’ base by breaking the non-covalent interaction between G1’ sugar pucker and pyrrole ring and at the same time rotate slightly on the plane perpendicular to the G-RNA stabilizing the mentioned π-π stacking interaction.

The available X-ray crystal structure ^65^ resolves two binding modes of TMPyP4: (a) top-face and (b) groove site interactions with specific bases such as C11 and C11’ bases. The results from WT-MetaD simulation complements the crystalline data and captures other binding possibilities of TMPyP4 to RNA G4 such as binding to C3’ base and also π-π interaction between pyrrole ring and sugar pucker ring, thereby highlighting the robustness of the present simulation. At the end of the simulation, basin M3 is found to be equally stable with the groove bound mode such as M1 and M2. Furthermore, the free energy difference between the top-face binding mode and groove binding mode is calculated to be ∼3.8 kcal/mol (ΔΔG_cal_) which is in good agreement with experimental ITC data (ΔΔG_expt_= 2.3 kcal/mol) (Table 1) The FES is able to identify a suitable pathway of binding/unbinding events of TMPyP4. The possible binding/unbinding events generally occur via the interaction between TMPyP4 and C11 nucleotide base resulting in the formation of the M1 basin (Figure 3). This particular base plays a key role in bringing TMPyP4 back to the top-face of the RNA G4 from bulk solvent. The rebinding mechanism of TMPyP4 to RNA G4 is shown in the Movie S1 in the Supplementary Information.

We then employed several orthogonal biophysical methods to further explore our computational findings. To study the conformational states of G4 structures formed from the RNA PQS18-1 sequence we employed circular dichroism (CD), a widely used analytical method that gives information about DNA structure and folding.^75^ The RNA PQS18-1 was first annealed in a buffer containing 10 mM lithium cacodylate (pH 7.0) with a metal ion concentration of 100 mM KCl.

The CD spectra of RNA PQS18-1 under these buffer conditions gave rise to a positive peak at 264 nm and a negative peak at 240 nm, which is consistent with a parallel-stranded RNA G4 topology (Figure 4). CD titration methods were then utilized to investigate the formation of the G4/ligand complex using TMPyP4.^76^ We observed that as TMPyP4 was added to the RNA PQS18-1, a concentration-dependent decrease in the CD signal at 264 nm was observed across all the concentrations, to a plateau at ∼60-70 eq. (60-70 μM). Meanwhile, although initially there was no signal observed at 295 nm, after addition of 1 eq of TMPyP4 (10 μM) a shoulder appears at this wavelength, consistent with remodeling of the RNA in an antiparallel formation during the binding process. As further equivalents of TMPyP4 are added, this signal increases up to 15 μM, reaches a plateau and then also decreases in a similar fashion to the parallel signal at 264 nm. Plotting the ellipticity at 264 nm against concentration of TMPyP4 added gave a sigmoidal shaped curve, indicating a co-operative process. We fitted this curve to the Hill equation, which identified Hill coefficients (n) of 4.1 ± 1.3. This reveals that the binding of TMPyP4 to PQS18-1 exhibits positive cooperativity (n > 1). The half degrading concentration ([DC]_50_) was determined to be 42 ± 0.5 μ M. These results are consistent with the unfolding mechanism observed in our modelling experiments. As a direct comparison we also performed a similar titration with the analogous DNA sequence using CD (Figure 5). In the DNA sequence equivalent, on addition of TMPyP4 there was no unfolding effect observed, however a prominent band at 445 nm appeared which increased in intensity with further additions of the ligand. We attribute this to an induced circular dichroism (ICD), indicative of strong binding between the DNA G4 and the TMPyP4. This direct comparison between both the DNA and RNA highlights the differences between the manner TMPyP4 interacts with G4 structures, depending on its composite nucleic acid.

**Figure 4.**
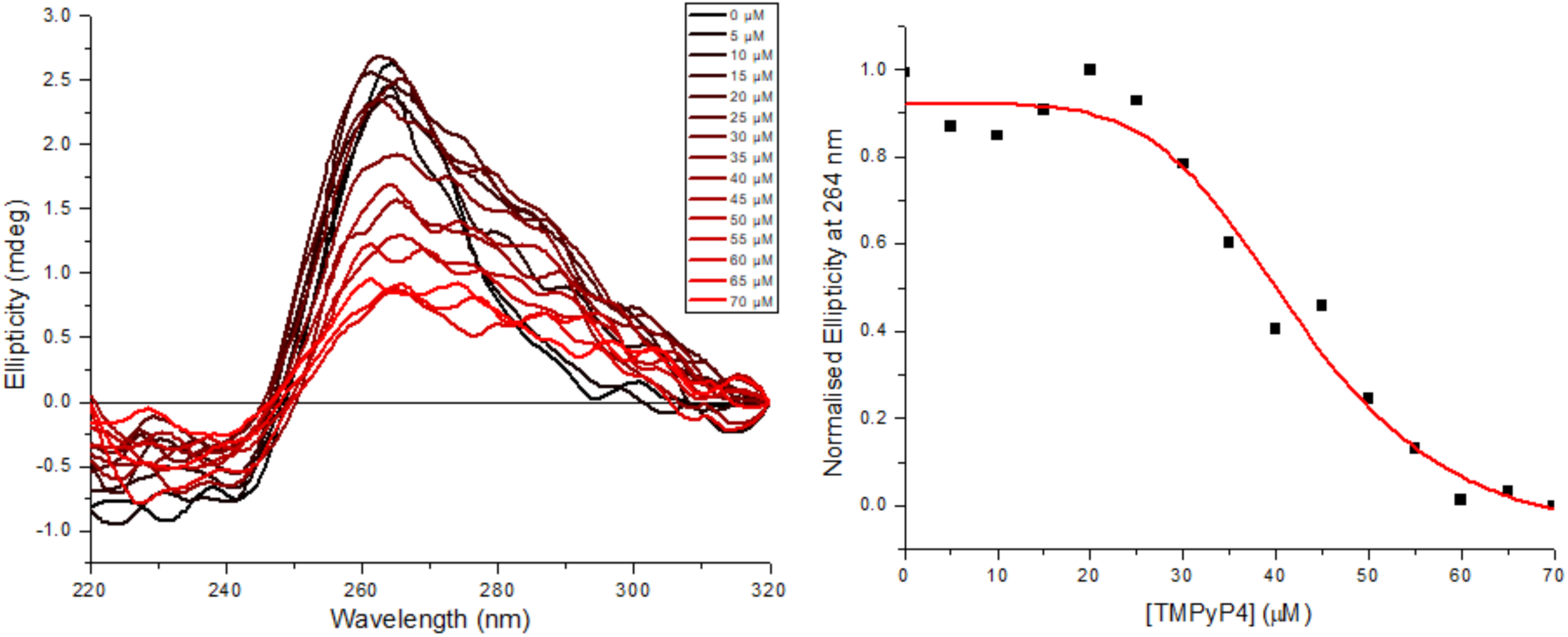
**Left:** CD titration of PQS18-1 RNA (10 µM) and TMPyP4 (0-70 µM) in 10 mM Lithium cacodylate 100 mM KCl, pH 7.0. **Right:** Plot of ellipticity at 264 nm against concentration of TMPyP4 and corresponding Hill fitting.

**Figure 5.**
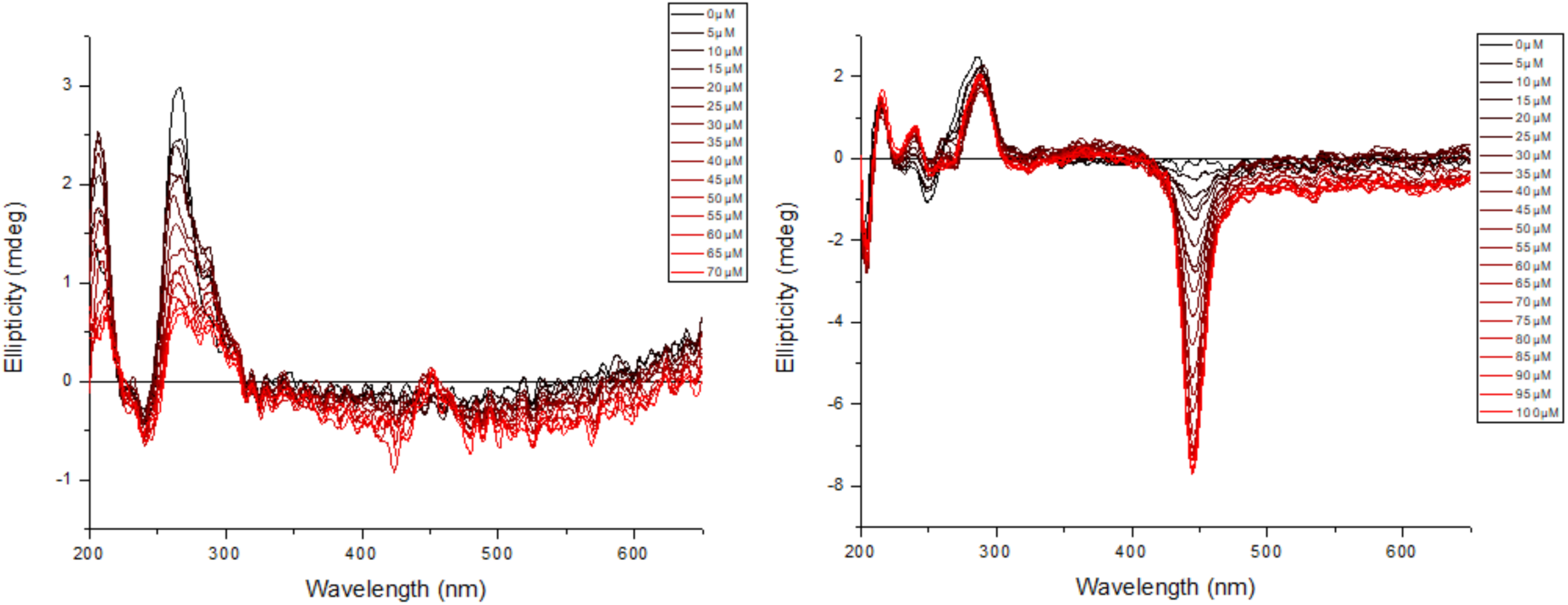
**Left** CD titration of PQS18-1 RNA (10 µM) in the presence of 0-70 μM TMPyP4. **Right** CD titration of PQS18-1DNA (20 µM) in the presence of 0-100 μM TMPyP4. Both experiments performed in 10 mM Lithium cacodylate 100 mM KCl, pH 7.0.

To further analyze the effect of TMPyP4 on RNA PQS18-1 we performed a Job’s analysis using CD to indicate the stoichiometry is between 1:1 and 2:1, averaging at 1.5:1 (Figure S8). We performed CD melting analysis to determine the ligand-induced effect of TMPyP4 on the stability of RNA PQS18-1. The melting of RNA PQS18-1 in the CD seemed to give rise to two melting events, one at 45°C and another at 73°C (Figure 6). On addition of 1 eq of TMPyP4 the shape of the curve starts to change but the overall ***T***_m_ values did not change. Addition of another equivalent of TMPyP4 changed the shape of the curve again, to give an overall ***T***_m_ of 71°C, which indicates a Δ***T***_m_ of -2°C. This change in the shape of the melting curve is further increased when the number of equivalents of TMPyP4 is increased to 5 eq where again there are two clear transitions at 37°C and 70°C, indicative of Δ***T***_m_ values of of -8°C and -3°C. These results indicate TMPyP4 has a destabilizing effect on the RNA G4 structure. After the melting experiments, we also studied the corresponding annealing experiments in the absence and presence of ligand. These indicated that the melting and annealing processes in the presence of TMPyP4 are not reversible, which is not unexpected for the liganded complexes (Figures S9-11). However, we did also observe precipitation of complex in the annealed samples, so the results from the annealing experiments are complicated by precipitation and aggregation processes. The corresponding UV-vis melting and annealing experiments were performed in parallel, but were also affected by TMPyP4 absorption in the same region as the DNA (Figures S12-15).

**Figure 6.**
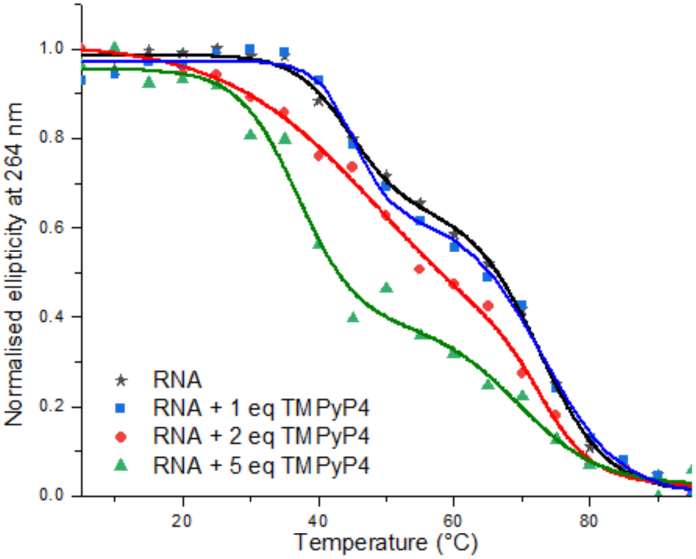
CD melting of PQS18-1 RNA (10 µM) in the presence of 0 eq (black stars), 1 eq (blue squares), 2 eq (red circles) and 5 eq (green triangles) of TMPyP4 in 10 mM Lithium cacodylate 100 mM KCl, pH 7.0.

We considered that some of the signal losses in the CD spectra might be through the effects of aggregation, rather than unfolding or disruption of the G4 structure, so during the CD titrations we also monitored the UV absorption spectra (Figure S16). Using this we were able to observe a linear dose-response on addition of TMPyP4 at 217 nm, consistent with Beer-Lambert law, indicating no aggregation at the concentrations examined at room temperature. Beyond these concentrations we sometimes observed precipitation, easily recognized as TMPyP4 is coloured. The apparently unfolding effects was observed at much lower concentrations than this and here we present data only at concentrations where precipitation was not evident.

UV-vis titrations of TMPyP4 with RNA titrated in (Figure 7) gave rise to a reduction and shift in the visible absorption spectra of TMPyP4 in the absence and presence of RNA PQS18-1 was observed. The strength and type of binding are indicated by the significant changes in wavelength maxima of absorption spectra upon addition of RNA G4, both the bathochromic shifts (Δλ = 20 nm) and hypochromicity (70%) shift to hyperchromicity (13%). Here the high values of bathochromic shifts and hypochromicity indicate strong stacking interactions between TMPyP4 and G-tetrads.

**Figure 7:**
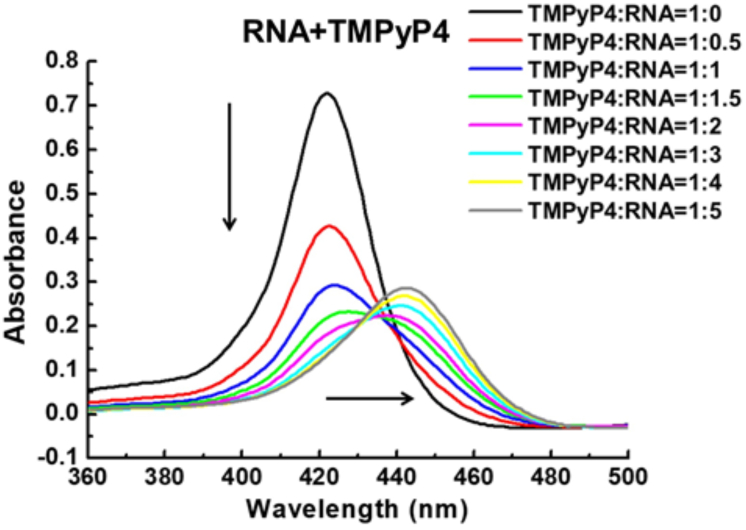
**UV titration spectra.** UV titration of TMPyP4 with RNA PQS18-1 in solution buffer (10 mM K_2_HPO_4_/KH_2_PO_4_ pH 7.0, 100 mM KCl). Conditions: [Ligand] = 4 μM; titrant: [RNA] = 0-20 μM. The vertical arrow indicates the decrease in the absorbance of TMPyP4 and the horizontal arrow indicates the shifted Soret band.

We also recorded UV spectra during the CD titrations between RNA PQS18-1 and TMPyP4 i.e. RNA with TMPyP4 added. Plotting the absorption at 440 nm against concentration gave a binding curve (Figures S17), which we fitted to a 2:1 binding model to indicate ***K***_d_s of 19 ± 0.7 μM and 188 ± 19 μM.

As an alternative to UV and CD, we also explored the apparent unfolding using FRET titrations, using dual labelled RNA PQS18-1, with FAM and TAMRA as a FRET pair. When in the folded G4 conformation the fluorophores will be in close proximity and FRET will occur, and it is possible to follow unfolding of G4 using this method. Folded RNA PQS18-1, when excited at 490 nm, gives rise to an emission spectrum with maxima at 515 nm and 585 nm, from the fluorophores FAM and TAMRA respectively. Addition of 0-5 eq of TMPyP4 to the folded RNA PQS18-1 caused a decrease in emission at 585 nm, indicative that TAMRA emission is decreasing, and potentially unfolding (Figure S18). TMPyP4 was also observed to cause an overall decrease in fluorescence emission through quenching, so we determined the FRET efficiency (***E***_FRET_) between the two fluorophores. A concentration-dependent decrease in ***E***_FRET_ was observed on addition of TMPyP4, consistent with unfolding of the RNA PQS18-1 structure in the presence of TMPyP4.

Finally, we next investigated the interaction between TMPyP4 and RNA PQS18-1 G4 by calculating complete thermodynamic parameters, including the stoichiometry and the values of free energy (ΔG), enthalpy (ΔH), and entropy (ΔS) changes of the binding reaction using Isothermal Titration Calorimetry (ITC). Correlating such thermodynamic data with a structural description increases our understanding of the molecular recognition process involved in complex formation and maintenance. The TMPyP4 (500 μM) was titrated into RNA solution (20 μM, 200 μl) with 10 mM K_2_HPO_4_/KH_2_PO_4_ pH 7.0, 100 mM KCl buffer at 25 °C. Figure 8 shows the integrated heat change data (after the correction of heat of dilution) and the corresponding heat changes are fit with multiple sites binding model. All the fitting parameters are summarized in Table 1. The two-site binding data was then fitted to a two-independent site binding model to determine the affinity and binding enthalpy of each of the two sites. Hence, the ITC experiments resulted in Ka_1_ value of (5.74 ± 0.46) × 10^5^ M^-1^, ΔH_1_ of -21.7 ± 0.9 kcal/mol, and ΔS_1_ of -46.5 cal/mol/deg (low affinity site), while the high affinity site has a Ka_2_ value of (2.83 ± 2.67) × 10^7^ M^-1^, a ΔH_2_ of -0.68 ± 0.4 kcal/mol, and a ΔS_2_ of 31.8 cal/mol/deg. The results indicated that the binding of TMPyP4 to two sites was different. The non-sigmoidal binding curve revealed that TMPyP4 could bind to the RNA at more than one site. The ITC data indicates that the stoichiometry of this multi-site interaction is two-site binding as the thermogram saturates after the addition of four molar equivalents of TMPyP4. The dip at the start of the ITC thermogram indicates that the second (lower affinity) site is more exothermic than the first (higher affinity) site (Figure 7). The enthalpy of the system starts decreasing as the second site starts to be populated. The total enthalpy is negative when both sites are saturated with the ligand. This data is strikingly correlated with our WT-MetaD simulation results where we found two stable binding sites as described above. The high and low affinity sites correspond to the top-face and groove-bound states, respectively. Since the groove bound state is completely solvent exposed as seen in the crystal data as well as from the WT-MetaD simulation, therefore it is evident that while moving from top-face bound state to groove bound state, the enthalpic contribution to the free energy of binding decreases as the solvation entropy plays a vital role in the ligand binding. The binding mechanism predicts that TMPyP4 is being caught by the fraying base C11 through π-π stacking interaction, where it forms the groove-bound state, showing the importance of the C11 base on the G4-TMPyP4 association. While sampling this particular state, the two top-face fraying bases A14 and A14’ remain in a wide-open conformational state due to the loss of H-bond between themselves, exposure to the bulk solvent and excessive flexible nature. Finally, TMPyP4 transfers from groove to the top-face of RNA and forms the top-face bound state by stacking on the terminal G-tetrad. This is followed by the loop closure and TMPyP4 is eventually sandwiched between the bases from terminal G-tetrad and top fraying bases A14 and A14 (for better understanding of the binding mechanism, see Movie S1 in Supplementary Information).

**Figure 8:**
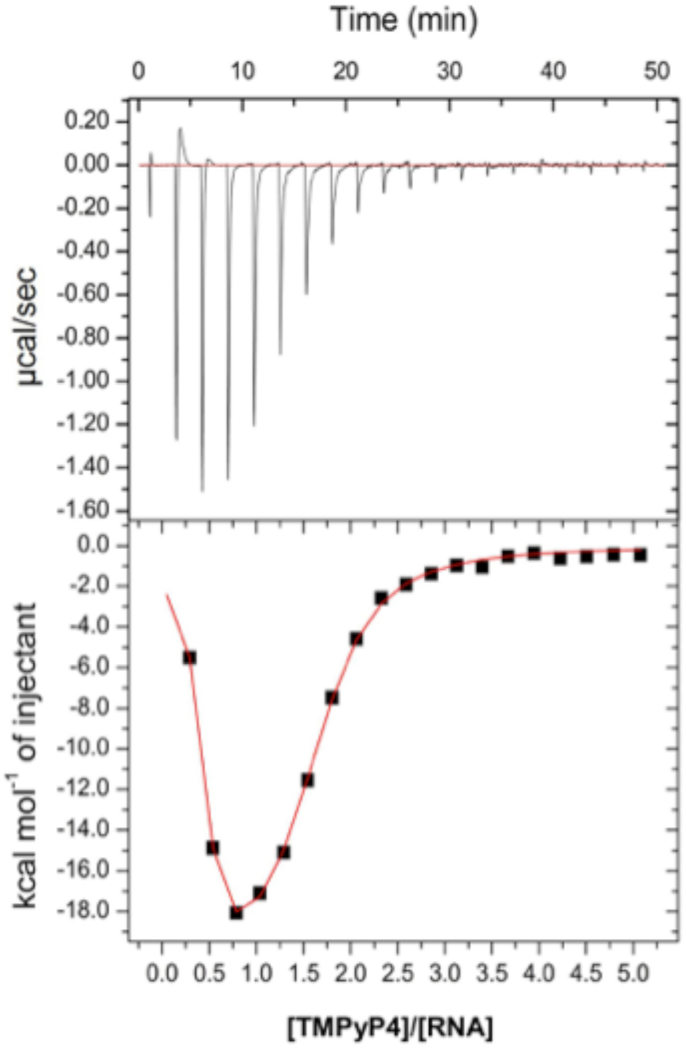
**ITC of TMPyP4-RNA G4 complex** ITC binding profile of titration of 500 μM TMPyP4 solution into 20 μM RNA PQS18-1 solution in 10 mM K_2_HPO_4_/KH_2_PO_4_ pH 7.0, 100 mM KCl buffer at 25 °C. Raw and fitted isotherms are shown in the panel. It is corrected for the corresponding heat of dilution by blank titration of titration of 500 μM TMPyP4 solution into buffer.

### The Unfolding Mechanism

To trace the unfolding mechanism of a bio-macromolecule using molecular simulation technique, such as MD simulation, one has to simulate the system sufficiently long since the sampling time for a typical unfolding remains on the scale of microsecond to millisecond.^77^ However, one can overcome this time scale problem using an enhanced sampling technique.^78, 79^ A combination of WT-MetaD and multiple walkers method has enabled us, for the first time, to characterize a possible TMPyP4 mediated unfolding mechanism of an RNA G4 topology.

In this report, we have identified two novel CVs, POA (position on the axis as d.z in nm) and DFA (distance from axis as R in nm), initially applied to study DNA ligand association process by O’Hagan et al.^74^ and adapted them to study RNA PQS18-1 G4 and its interaction with TMPyP4. These CVs can configure and describe all the possible bound/unbound states along with unfolded states. The FES highlights the connections between the described states (Figure 9a). The groove-bound state, top-face and unfolded states are assigned with the following distance in the d.z versus R curve: -1.5 ≤ d.z ≤ 0.7 nm and 1.0 ≤ R ≤ 2.0 nm; 0.8 ≤ d.z ≤ 1.5 nm and 0.0 ≤ R ≤ 0.5 nm; and -0.5 ≤ d.z ≤ 0.2 nm and 0.0 ≤ R ≤ 0.5 nm, respectively. A cartoon representation on the possible mechanism of the TMPyP4 mediated unfolding is illustrated in Figure 9c. As observed in the FES, the unfolding region is close to ∼0.0 nm, suggests that the unfolding occurs via the intercalation process as the CV distance between the DNA and ligand decreases. In the first step, C11 base pulls TMPyP4 back on the top-face of the RNA i.e. the rapid transfer of ligand occurs from groove-bound state to the top-face (see T1 and T2 in Figure 2 and Supplementary Information, Movie S1, where the ligand rebinding process is illustrated). Before TMPyP4 jumps toward top-face bound state, the groove bound-state remains in a closed-like conformation (Figure S3). As TMPyP4 moves to the top-face, the RNA conformation shifts from a closed to an open state. This open conformation of the RNA is also observed in our unbiased MD simulation and resembles the top-face bound state (Figure S3). In the second step, TMPyP4 slides down towards the groove, stays in a slight tilted conformation and interacts with the bases from the first G-tetrad such as G10 and G13’. In the next step, TMPyP4 intercalates through the first and second G-tetrad by breaking their associated Hoogsteen-bonds. As a result, TMPyP4 finally reaches the third G-tetrad by breaking the RNA G4 topology completely to an unwound state (Supplementary Information, Movie S2).

**Figure 9:**
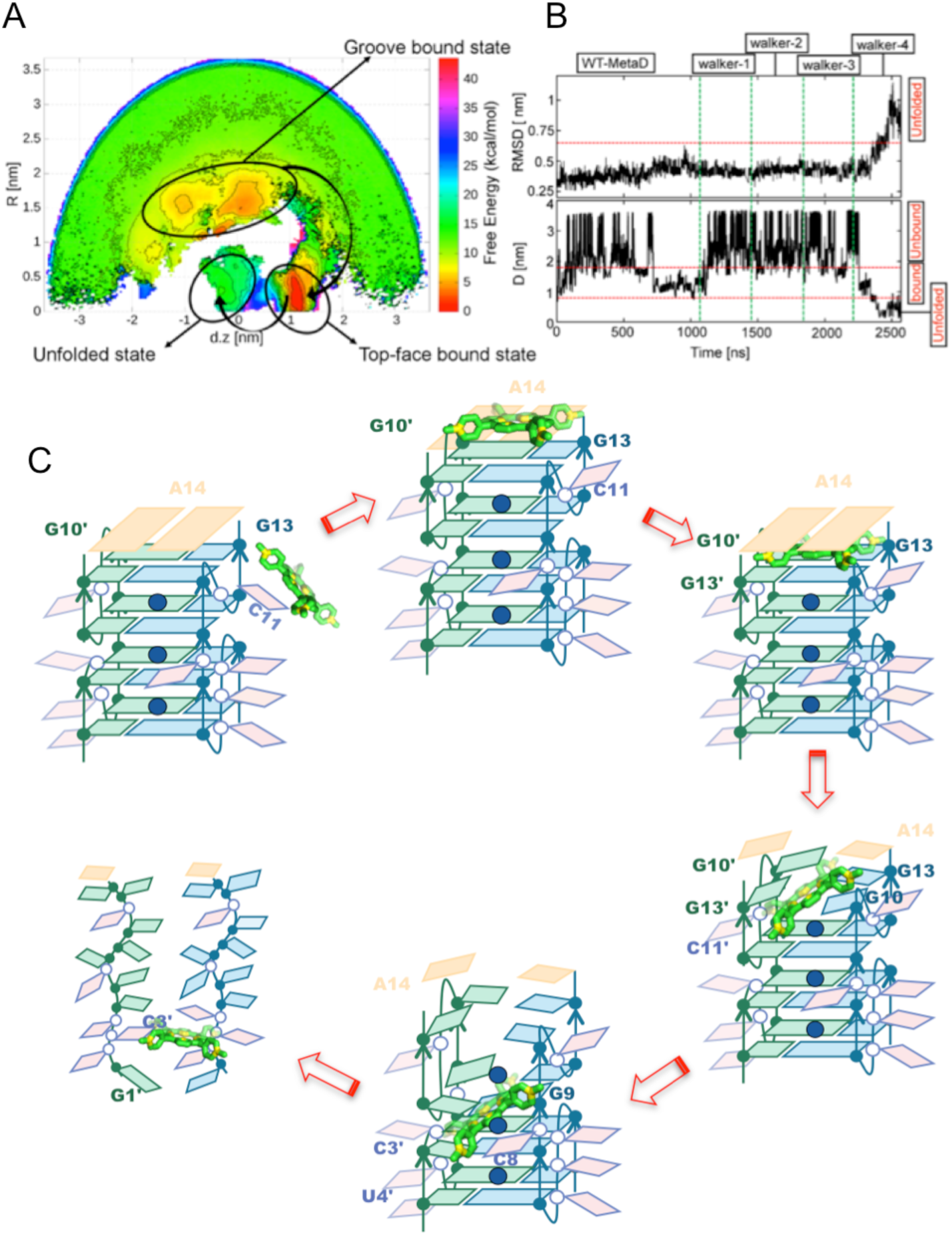
**The unfolding mechanism.** (a) Two-dimensional free energy surface plotted as a function of POA (position on the axis as d.z in nm) and DFA (distance from axis as R in nm), highlighting the pathway between various states. TMPyP4 mediated unfolding is initiated via stacking interactions identified in the top-face bound state. (b) The RMSD of the RNA G4 backbone throughout the 2.6 µs of simulation time (top). As observed in the last replica (walker-4), RMSD increased from an average value of ∼3.5 nm to > 1.0 nm (thereby decreasing the CV distance from being ∼1.0 nm i.e., from the top-face bound state to ∼0.2 nm) indicating unfolding via intercalation. The simulation statistics, illustrating the distance between TMPyP4 and RNA G4 where all the walkers are combined together (bottom). (c) Proposed model for ion-assisted recognition and unfolding mechanism of G4 by TMPyP4. Guanine nucleotides are in blue (chain A) and green (chain B). Loop nucleotides are in pink. Potassium ions are in blue. TMPyP4 is illustrated in green sticks with nitrogen atoms are in yellow.

## Discussion

The present work reflects on the complexity of ligand interactions on the stability of the G4s. We show that the ligands could behave both as stabilizers or destabilizers of G4s based on how they interact with the G4s. TMPyP4, has been reported to both, stabilize and unfold G4s.^42, 45, 49, 80^ The scarcity of the dynamic structural information about the ligand binding makes it necessary to rely heavily on biophysical characterization. Job plot indicated that the stoichiometry of TMPyP4 binding to PQS18-1 RNA G4 is more than 1:1, indicating binding at two or more distinct sites or modes (Figure S9). The “two independent sites” model exhibited in the ITC assay showed that more than two states exist in the unfolding processes (Figure 8). When ligand:DNA = 3:4 (ITC experiment)/ 2:3 (UV experiment), both the UV absorption and ITC thermogram exhibited a big change. This concentration-dependent biophysical character variance suggests the existence of a structural intermediate in the titration process. Importantly, our biophysical experiments were performed at a range of concentrations of RNA, from 0.1 μM (in the FRET titrations) to 20 μM (in the ITC experiments). This indicates the unfolding effect is observed at a wide range of concentrations of RNA, and is not limited to when the RNA is at higher concentrations.

The ITC experiments show that TMPyP4 could completely disrupt the RNA G4 structure. This extent of TMPyP4 disruption was plausibly due to the higher melting temperature of the RNA PQS18-1 G4 structure (***T***_m_=68.4°C). A lower concentration of TMPyP4 was required for RNA PQS18-1 destabilization at higher temperatures. We found that the apparent initial rates of G4 unfolding decreased as a function of thermal stability imparted by ligand binding. While previous reports have indeed shown that high potassium concentrations have a stabilizing effect on the ***T***_m_ of various G4s, it is not trivial to conclude that the unfolding kinetics positively correlates with the thermal stability.

Based on our simulation and experimental results we propose that the observed unfolding could be broadly divided into three steps: (a) TMPyP4 approaches PQS18-1 RNA G4 and forms an initial complex structure; (b) The stability of the intermediate complex depends upon the dynamic interactions between the ligand and the G4 structure. The dynamic interactions determine whether small molecules stabilize or unfold a G4 structure and (c) the strength of the dynamic interactions between the ligand and G4 structure determines the fate of the inter-quartet cations. In the case of destabilizers, like TMPyP4, the inter-quartet cations are expected to be ejected, which accelerates the unfolding process.

The binding mode of TMPyP4 to G4 remains controversial: some reports say that it binds and stabilizes G4s,^57, 61^ a few suggest that TMPyP4 alters the G4 structure,^80^ while others have demonstrated that it unfolds G4s.^41, 45, 49^ Here we show that all types of binding coexist, and that the distribution of final states depends on the ligand concentration ratio and the temperature. Finally, it is becoming increasingly clear that it is extremely difficult to predict the behavior of G4 interacting ligands on different G4 topologies and nucleic acids. This emphasizes that ligands cannot be designed based on the traditional approaches of exploiting the topology, without taking dynamic interactions into account.

## Methods

### Standard (Unbiased) MD simulations

We carried out standard (unbiased) MD simulations of all the available TMPyP4 bound poses to RNA G4. Initial crystal structure revealed a top-face bound state of TMPyP4 where it is sandwiched in between the top fraying base A14 and A14’ and the bases G10, G10’, G13 and G13’ from the first G-tetrad. In the second bound state TMPyP4 stack on the C11 base and it is mainly found to be a solvent exposed (Figure 1).

All the unbiased simulations were performed in triplicates using the Gromacs-5.0 software package.^82^ The bsc0χ_OL3_ force field was used for the RNA parameterization.^84–86^ For the ligand, the General Amber Force Field (GAFF) were used to generate parameters.^87^ The charges were calculated using the restrained electrostatic potential (RESP) fitting procedure.^88, 89^ The RESP fit was performed onto a grid of electrostatic potential points calculated at the HF/6-31G(d) level as recommended by many works.^90, 91^

The K+ ion in the central axis of the structure were treated as an integral part of the structure. The RNA-TMPyP4 complex was solvated in a cubic box with the dimension of 7.8 x 7.8 x 7.8 nm^3^ along with 15364 TIP3P explicit water molecules.^92^ The Joung and Cheatham cations optimized for TIP3P water were used to generate 100 mM KCl concentration of the system.^93^ The RNA-TMPyP4 complexes were minimized before the equilibration and production run as follows: the minimization of the solute hydrogen atoms on the RNA and TMPyP4 was followed by the minimization of the counterions and the water molecules within the box. In the next step, the RNA backbone along with the all the heavy atoms on TMPyP4 were restrained, and the solvent molecules with counter ions were allowed to move during a short 50 ps MD run, therefore, relaxing the density of the whole system. In the next step, the nucleobases were relaxed in several minimization runs with decreasing force constants applied to the RNA backbone atoms, however, only few phosphate atoms were kept restrained with a force constant of 0.239 kcal/mol/nm^2^. After the full relaxation, the system was slowly heated to 300K using velocity rescaling thermostat ^94^ with a coupling constant of 0.5 ps employing NVT ensemble. As the system reached to the temperature of interest (300 K), the equilibration simulation was performed for 10 ns using an NPT ensemble with Berendsen thermostat and Berendsen barostat,^95^ and 0.5 ps was used again as the coupling constant for both temperature and pressure, respectively. Finally, the production run was set for 1 μs using Nose-Hoover thermostat ^96^ and Parrinello-Rahman barostat ^97^ with the same coupling constant as previously taken in the equilibration simulation in the NPT ensemble. All the simulations were carried out under the periodic boundary conditions (PBC). The particle-mesh Ewald (PME) method was used to calculate the electrostatic interactions within a cut-off of 10 Å.^98, 99^ The same cut-off was used for Lennard-Jones (LJ) interactions. All simulations were performed with a 1.0 fs integration time step.^97,96,97^

### Well-tempered Metadynamics simulation

We performed a well-tempered variant of the metadynamics simulation of TMPyP4 binding to RNA G-quadruplex. The WT-MetaD helps to understand a complete binding/unbinding mechanism of TMPyP4. Further, a pathway for the TmPyP4 mediated unfolding of the RNA G4 is disclosed for the very first time. The WT-MetaD simulation was started with a well-equilibrated structure generated from the crystal structure of the RNA-TMPyP4 top-face bound state. In particular, the starting structure for the WT-MetaD simulation was taken after 20 ns of unbiased MD simulation, which was found to be a rather stable ligand-binding conformation. We performed over 2.5 μs of WT-MetaD simulation ^100^ in order to obtain an accurate estimate of RNA-TMPyP4 binding free energies through sampling all the individual states such as bound, unbound and unfolded states. We used a combination of the following scheme:

1. Well-tempered variant (WT) of the metadynamics
2. The multiple walker technique, placing four walkers based on the TMPyp4 binding sites:

a. Top-face bound state (one-replica)
b. Groove bound state (two-replica)
c. Unbound state (one-replica)

The same systems and MD settings as described previously were used for the WT-MetaD simulation. The plumed 2.3 plugin ^101, 102^ was used to carry out the simulation with the Gromacs-5.0.7 code.^82^ The bias potential was calculated according to the WT-MetaD scheme as follows:

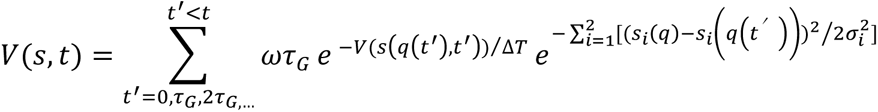

where the deposition rate, ω, and deposition stride, τ_G_, of the Gaussian hills were set to 0.358 kcal/mol/ps (1.5 kJ/mol/ps) and 1.0 ps, respectively. The bias factor (T + ΔT)/T was set to 15, and the final FES was calculated as follows:

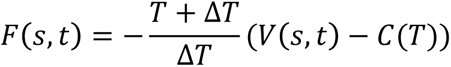

where the V(s,t) is the bias potential added to the Collective Variables (CV) used and the T represents the simulation temperature. ΔT is the difference between the temperature of the CV and the simulation temperature. The bias potential is grown as the sum of the Gaussian hills deposited along the chosen CV space and finally the sampling of particular CV space can be controlled with the tuning of the ΔT parameter.^66^ The torsions used in the metadynamics simulations are illustrated in Figure S19.

### Analysis

As discussed in the previous section, the X-ray crystal structures ^103^ were downloaded from the PDB data bank and prepared for the MD simulation. The visualization and analysis of the MD statistics are performed through VMD molecular visualization program.^104^ Hydrogen bond analysis of the quartets (Figure S20) was carried out using analysis tools implemented in the VMD. All the necessary graphs are first prepared with Gnuplot program (http://gnuplot.info) and later modified with Gimp 2.0 (https://www.gimp.org) program. The analysis of the Metadynamics simulation is performed through the Plumed 2.3 package.^102^ The high resolution figures are prepared through PyMOL program (www.schrodinger.com).

### Materials

Experiments based in China (UV and ITC) were performed using oligonucleotide sequence RNA PQS18-1 5ʹ-[r(GGCUCGGCGGCGGA)]-3ʹ purchased from Tsingke biological technology (Beijing, China) and TMPyP4 were purchased from Frontier Scientific (Logan, Utah, USA) and Sigma Aldrich. The concentration was determined using the Beer–Lambert law by measuring the absorbance at 260 nm using Nanodrop Photometer N60 (Implen, Germany). The extinction coefficients were obtained from the IDT Web site (https://sg.idtdna.com/calc/analyzer). Further dilutions were carried out in buffer containing 10 mM K_2_HPO_4_/KH_2_PO_4_ pH 7.0, 100 mM KCl. The starting oligonucleotide solution were annealed in the corresponding buffer by heating to 95 °C for 5 min, followed by gradual cooling to room temperature.

Experiments based in the UK (CD, UV and FRET) were performed using oligonucleotides purchased from Eurogentec (PQS18-1 5′-[r(GGCUCGGCGGCGGA)]- 3 ′ PQS18-1DNA 5 ′ -[d(GGCTCGGCGGCGGA)]-3 ′ and PQS18-1FRET 5 ′ -[FAM-r(GGCUCGGCGGCGGA)-TAMRA]-3′), each of them were purified using reverse phase HPLC and supplied dry. The dry RNA/DNA was dissolved in nuclease-free water in order to preTpare stock solution, the FRET-labelled RNA sample weas prepared at approximately 100 µM and the unlabelled RNA prepared at 1 mM. The concentration of the stock solutions was identified from their UV absorbance at 260 nm with a NanoDrop by using extinction coefficients that provided by Eurogentec, further dilutions were carried out in buffer containing 10 mM lithium cacodylate pH 7.0, 100 mM KCl. RNA and DNA samples were annealed using a heating block, samples were put into the block for 95 °C for 5 minutes followed by gradual cooling to room temperature and left overnight.

### Circular dichroism (CD) and UV-vis spectroscopy

CD experiments were performed with the JASCO 1500 spectropolarimeter (JASCO, Japan) under a constant flow of nitrogen. All measurements were done at 25°C with 1 mm quartz cuvette and covering a spectral range of 200–650 nm using a scan rate of 200 nm/min and with a 1 s response time, and 1 nm bandwidth. Each spectrum was obtained as an average of 4 measurements and was buffer/ligand subtracted, zero-corrected at 320 nm and smoothed using a Savitsky-Golay 5 point window. During titrations with TMPyP4 spectra were taken immediately after addition of TMPyP4. Unfolding data was fitted using the Hill equation θ = *C*^n^/(DC_50_^n^+*C*^n^) where θ = fraction of bound TMPyP4, ***C*** = concentration of TMPyP4, n = Hill coefficient, DC_50_ = the half degrading concentration.

Corresponding UV-vis absorption data was also collected at the same time as the CD. UV-vis binding data was fitted to a two inequivalent sites binding model: θ = (***K***_1_***C***+ 2***K***_1_***K***_2_***C^2^***)/(1 + ***K***_1_***C*** + 2***K***_1_***K***_2_***C^2^***) where θ = fraction of bound TMPyP4, ***C*** = concentration of TMPyP4, ***K***_1_ and ***K***_2_ are the association constants for the first and second sites.

CD/UV-vis melting/annealing profiles were recorded by monitoring the CD at 264 nm, as a function of temperature using a sealed cuvette. RNA samples were cooled to 5°C and then heated to 95°C with a heating rate of 1°C/min held at 95°C for 5 minutes and then cooled to 5°C at the same rate. The ***T***_m_ values of the complexes were calculated by normalizing the experimental curves to give fraction folded and the data fitted to sigmoidal fittings to determine the ***T***_m_ values. Final data was analysed in OriginPro 2020.

### UV-vis spectroscopy

Initially, 150 μL solutions of the blank buffer and the ligand sample (4 μM) were placed in the reference and sample cuvettes (1.0 cm path length), respectively, and then the first spectrum was recorded in the range of 200–600 nm. During the titration, an aliquot of buffered RNA solution was added to cuvette. Complex solutions were incubated for 5 min before absorption spectra were recorded. The absorption spectra were recorded on a UV1800 spectrophotometer (Shimadzu Technologies, Japan) at 25 °C.

### Fluorescence (FRET) Experiments

FRET titration experiments were performed in Edinburgh Instruments FS5 Spectrofluorometer, using a quartz cuvette with 10 mm path length. The sample volume was 250 µL. The FRET labelled RNA was diluted in pH 7.0 buffer to 0.1 µM. the sample was excited at 490 nm and the fluorescence emission was measured from 500 nm to 650 nm. TMPyP4 was diluted in buffer and added into sample in 0.025 µM increments from 0.025 µM to 0.5 µM. The relative FRET efficiency (***E***_FRET_) was calculated using ***E***_FRET_ = (***I***_a_/***I***_d_+***I***_a_ ) where ***I***_d_ is the fluorescence intensity of the donor and ***I***_a_ is the fluorescence intensity of the acceptor. The experiment was performed in triplicate and the error bars represent the standard deviation. The data was analysed by OriginPro 2020.

### Isothermal titration calorimetry (ITC)

ITC experiments were performed on a Auto-iTC100 titration calorimetry (MicroCal) at 25 °C. All solutions (buffer, RNA and ligand) were degassed before carrying out the experiments. A 20 μM solution of RNA PQS18-1 was placed into the cell and 500 μM TMPyP4 were taken in the rotating syringe (750 rpm). A total of 40 μL TMPyP4 was added to the RNA in 20 injections and the time gap for two injections was 150 s, while the first injection was 0.4 μL to account for diffusion from the syringe into the cell during equilibration. This initial injection was not used in fitting the data. Similarly, dilution experiments were also carried out taking buffer (10 mM K_2_HPO_4_/KH_2_PO_4_ pH 7.0, 100mM KCl) in the cell and TMPyP4 was kept in the syringe. We have used Microcal ITC Analysis Software for the analysis of the ITC raw data using two site-binding model. We have subtracted the dilution data from the raw data of the interaction of TMPyP4 with RNA before analysis. After fitting these experimental data, the enthalpy change during the process was obtained.

## Supporting Information

PC analysis of bound states, native RNA, top-face RNA-TMPyP4 bound statem groove site RNA-TMPyP4 bound state; description of the free energy surface; circular dichroism melting; 1D PMF; FES of native RNA G4 topology, top-face TMPyP4-RNA, groove bound TMPyP4-RNA complex, most populated clusters of RNA-TMPyP4 complex; Construction of CVs; MD simulation statistics; Job plot; CD melting/annealing; UV melting/annealing; UV-vis titration; Fluorescence titrations; details of CVs; movies.

## Data Availability

The unbiased simulation trajectories have been uploaded to zenodo (doi 10.5281/zenodo.5594466). Metadynamics simulation trajectories can be obtained from the authors upon request.

## Acknowledgements

DGW is supported by the National Natural Science Foundation of China (21732002, 22077043, 31672558). AJM and Susanta Haldar would like to acknowledge EPSRC grant for this study (EP/N024117/1 and EP/M022609/1). SH would like to thank Dr Gary N Parkinson for critical reading of the manuscript.

