## Supplementary file for "Ligand-induced unfolding mechanism of an RNA G-quadruplex"

| | **Contents** | **Page Number** |
|---|---|---|


| | | |
|---|---|---|
| | PC Analysis of the bound states | 4 |
| | PC Analysis on the native RNA | 4 |
| | PC Analysis on the top-face RNA-TMPyP4 bound state | 6 |
| | PC Analysis on the groove site RNA-TMPyP4 bound state | 7 |
| | Description of the free energy surface | 9 |
| Figure S1 | 1D Potential of mean force projected on distance, CV of the binding of TMPyP3 to RNA G4 | 10 |
| Figure S2 | FES showing the backbone fluctuation of native RNA G4 topology | 11 |
| Figure S3 | FES showing the backbone fluctuations of top-face TMPyP4-RNA complex | 12 |
| Figure S4 | FES showing the backbone fluctuations of groove bound TMPyP4-RNA complex | 13 |
| Figure S5 | The FES (D vs T) showing the most populated clusters of RNA-TMPyP4 complex on the groove binding position | 14 |
| Figure S6 | Construction of CVs | 15 |
| Figure S7 | MD simulation statistics of TMPyP4-groove bound states | 16 |
| Figure S8 | Job Plot of RNA PQS 18-1 and TMPyP4 | 17 |
| Figure S9 | CD melting and annealing of RNA PQS18-1 | 17 |
| Figure S10 | CD melting and annealing of RNA PQS18-1 and 2 eq TMPyP4 | 18 |
| Figure S11 | CD melting and annealing of RNA PQS18-1 and 5 eq TMPyP4 | 19 |
| Figure S12 | UV melting and annealing of RNA PQS18-1 | 20 |
| Figure S13 | UV melting and annealing of RNA PQS18-1 + TMPyP4 (2 eq) | 20 |

| | | |
|---|---|---|
| Figure S14 | UV melting and annealing of RNA PQS18-1 + TMPyP4 (5 eq) | 21 |
| Figure S15 | UV melting and annealing of RNA PQS18-1 in the absence/presence of TMPyP4 | 21 |
| Figure S16 | UV titration spectra and plot of UV absorbance | 22 |
| Figure S17 | UV absorbance at 440 nm | 22 |
| Figure S18 | Fluorescence spectra of dual labelleled RNA PQS18-1 | 23 |
| Figure S19 | Torsion definition | 24 |
| Table S1 | List of atoms with parameters that are involved in building the distance and torsion CVs | 24 |
| Figure S20 | N1-O6 and N2-N7 distances in quartets | 25 |
| Movie S1 | The dynamics and rebinding mechanism of TMPyP4 to RNA G4 | 26 |
| Movie S2 | The RNA unfolding mechanism by TMPyP4 | 26 |

### Supplementary Discussion

#### PC Analysis of the bound states

To get an insight into the dynamical behaviour of the RNA itself and further to understand the unfolding pathway found in the metadynamics simulation, we performed Principal Component Analysis (PCA) to each of the TMPyP4 bound states from the unbiased MD simulation. The three most stable complexes between RNA and TMPyP4 are shown in Figure 1, 2 and 3 in the main text. The most stable basin is the top-face bound state where TMPyP4 interacts with A14, A14' and G10, G10', G13 and G13' bases. The next stable basins following the top-face one belong to the major groove bound states where TMPyP4 interact individually with C11 and C11' bases. We performed ~1 μs (800 ns) standard MD simulation to each of these binding poses to study their dynamics.

#### PC Analysis on the native RNA

PCA calculations were performed using Gromacs software package using gmx cluster facility; version-5.0.7. The starting structure of RNA G4 for the MD simulation is taken from the X-ray crystal structure which shows an open-like backbone conformation. Only the RNA G4 (without TMPyP4) was used for PCA analysis. The PCA analysis on the native RNA after 800 ns of standard unbiased MD simulation is shown in Figure S2. The FES obtained using PCA1 and PCA2 revealed four stable basins which correspond to the changes in the conformations of the RNA backbone. A cluster analysis was performed using only the backbone heavy atoms and the most populated cluster from each basin was obtained. Basin A depicts a X-ray crystal-like structure (PDB id: 6JJH) where both the top Adenine bases, A14 and A14', stacked on the G13 and G13' bases of the first G-tetrad. Some fraying behaviour of A14 and A14' was also observed. In particular, A14 base is flipped over G13 base which disrupts the stable H-bond between both the Adenine bases allowing A14 base to fray towards solvent. At the bottom of G-tetrad a crystal-like open conformation of the backbone was observed where capped bases such as C5 and C5' bases are engaged in H-bond formation while stacking on the G6' and G1 bases of the last/bottom G-tetrad, respectively (see Figure 1 from main text). In the next basin, basin B, where the backbone containing A14 base is flipped and partially stacked with G13 base from the groove site (Figure S2). It is worth mentioning that such conformational change is also seen in the metadynamics simulation when TMPyP4 leave the top-face bound state for solvated state. Looking at the bottom site, one can again see an open like conformation where C5 makes a π–π stacking interaction with C3 base thereby increasing the H-bond distance from 2.8 Å to 3.7 Å between C5 and C5'base. In the open like conformation, both the C5 and C5' bases are intact with the formation of a H-bond interaction and with the increase or decrease of this H-bond distance, the RNA backbone goes far or come close to each other, respectively. In basin C, a large backbone conformational change is seen. Especially, two specific non-covalent interaction is formed: a H-bond between C5' base and the oxygen atom of the backbone and a π-π stacking between C5 and C5' base. Since the specific H-bond between C5 and C5' base is now broken, and also due to the newly formed π–π stacking interaction, both the bases come close to each other bringing chain A and chain B close and forming the close like conformation. Such conformational change provides evidence of the native RNA being flexible over a short period of time. In the basin D, the conformation is more or less similar to the basin C in which C3 bases come closer to the C5' base and formed a stable H-bond network. Moreover, the top-face conformation does not change throughout the simulation. Overall, the PCA analysis revealed the flexibility of RNA structure and hence it can adapt to a range of conformations.

**PC Analysis on the Top-face RNA-TMPyP4 bound state**

The PCA analysis of the RNA-TMPyP4 top-face bound state is performed after ~1 μs (800 ns) MD simulation. The starting structure of the RNA G4 TMPyP4 complex for the MD simulation is again taken from the X-ray crystal structure (PDB id : 6JJH) which shows an open-like backbone conformation. The two-dimensional FES using PCA1 and PCA2 reveals three most stable basins (Figure S3). Basin A represents the starting conformation where the RNA backbone retains its open like conformation which is similar to the conformation found in basin A for native RNA PCA analysis (Figure S2). At the bottom, one can see that an open-like conformation exists following the two typical non-covalent interaction: a H-bond between C5 and C5' base and a π–π stacking interaction between the C5' and C3' base. TMPyP4 is bound on top of the RNA being sandwiched in between the fraying bases A14, A14' and G10, G10', G13 and G13' bases from the first G-tetrad employing π–π stacking interactions. In basin B, TMPyP4 maintained the same conformation which suggests that the ligand stabilizes the top-face loop fluctuation incorporating A14 and A14 bases relative to native RNA (Figure S1 for the corresponding loop fluctuation in the absence of TMPyP4). At the bottom, the backbone still maintains an open-like conformation as basin A. The only difference is found to be a slight increase in the H-bond distance between C5 and C5' base which helps to maintain open backbone conformation between two single RNA strands (chain A and chain B). In basin C, the top-face is again seen to be stable without any significant change in the structure relative to basin A and B. However, looking at the bottom, a notable difference can be seen in terms of the absolute position of the bases. In particular, the U4 base flips from its fraying position in basin B to the $\pi$–$\pi$ stacking conformation with the G1 base allowing the C5 base to shift its position from being stacking with G1 base to G1' base. These rather large conformational shifts by both U4 and C5 bases disrupt the $\pi$–$\pi$ stacking interaction between C5' and C3' base, thereby allowing the backbone to shift back and finally maintain an open-like conformation again. Overall, the PCA analysis on the ~1 μs MD simulation revealed that RNA was stabilized by the complex formation with TMPyP4 which in particular do not allow large conformational changes. Moreover, RNA was maintained in an open-like conformation throughout the simulation.

**PC Analysis on the Groove site RNA-TMPyP4 bound state**

We have performed the PCA analysis on the RNA-TMPyP4 groove site binding state to compare with the native and top-face complexed state. Specifically, C11 (RNA)…TMPyP4 groove bound state has been thoroughly analysed. The FES obtained after ~1μs (800 ns) unbiased MD simulation employing PCA1 and PCA2 reveals three most stable basins (Figure S4). A cluster analysis was carried out as earlier using the backbone heavy atoms to obtain the representative structures of the basins. Finally, the populated clusters are thoroughly represented along the FES from each basin. Basin A represent a rather close-like conformation where both the C5 and C5' bases are neither engaged in the H-bond formation nor they are in the $\pi$–$\pi$

stacking interaction between them; rather they are engaged in the H-bond formation with backbone oxygen atoms from opposite RNA chains while the ligand is bound to the C11 base. Additionally, C5' base forms an extra H-bond with C3 base. These large H-bond networks bring both the RNA strand close to each other resulting in a close-like conformation. Such a typical H-bond network is already seen in the native RNA PCA analysis (see basin D, Figure S2). In basin B, one can see a slight change in the conformation of the C3 base and thus the backbone of overall RNA. Specifically, the C3 base flips over the G1 base which disrupts the H-bond between C5' and C3 base but the RNA still remains in its close-like conformation. In basin C, a rather large shift in some specific bases were observed compared to the overall conformational change in the backbone. The H-bond network in the top-face was broken due to a large conformational shift of the A14' base, however at the same time, A14 base maintained its original position by stacking on the G13 base of the first G-tetrad. These findings suggests that the top-face loop is stabilized only by the formation of RNA-TMPyP4 top-face complex. Further, at the bottom, both the C5 and C5' bases are engaged in the $\pi$–$\pi$ stacking interaction maintaining the overall close-like conformation. Such a conformation again resembles with one of the native RNA conformations found earlier (see basin C and D, Figure S2). Overall, it is worth mentioning that the starting structure for the RNA-TMPyP4 groove-bound complex, taken from the WT-MetaD simulation, is a close-like conformational state which stabilizes the RNA by the complex formation and maintains its conformation throughout the simulation. Moreover, it is also evident that TMPyP4 groove-bound states always maintain a close-like conformation which is opposite to the top-face bound states.

**Description of the Free Energy Surface**

The D vs T FES does not separate the groove binding sites with top-face ones (Figure S5). Three most populated clusters are shown which represent the basin M1 in the groove position of the FES ranges from 1.4 nm to 2.1 nm as shown in Figure 1 (see main text). Cluster 1 represents the RNA-TMPyP4 groove binding mode where TMPyP4 makes π–π stacking with C11 base from chain A. In particular, the indole rings from the porphyrin ring in TMPyP4 is taking part in the π–π stacking with C11 base from RNA. Cluster 2 appear for an RNA-TMPyP4 complex with top-face binding conformation. Here the ligand is stacking on the top-face of the RNA interacting with fraying adenine base such as A14' and partially with A14. Cluster 3 constitute for another RNA-TMPyP4 groove binding conformation where again a π–π stacking is seen in between C11' base from chain B and TMPyP4.

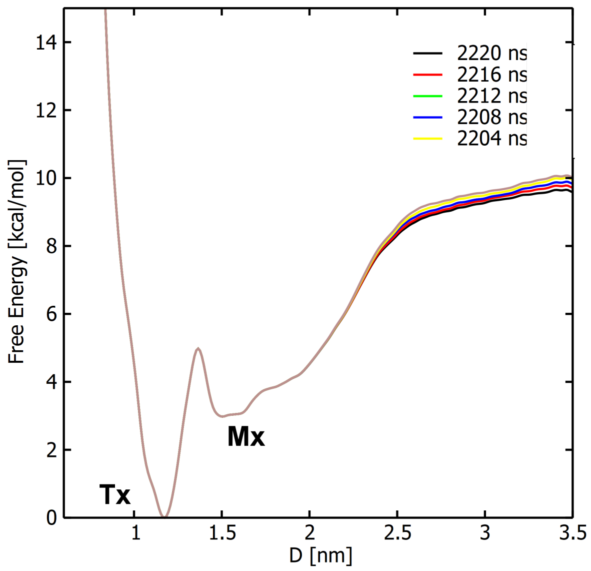


**Figure S1: The one-dimensional potential of mean force (PMF) projected on the distance (D), CV of the binding of TMPyP4 to RNA G4**. The free energy profile is monitored in the last 16 ns going from 2204 to 2220 ns showing the convergence of the free energy with the difference in the free energy being <0.4 kcal/mol. The last replica (walker-4) is ignored due to sampling of the unfolded region.

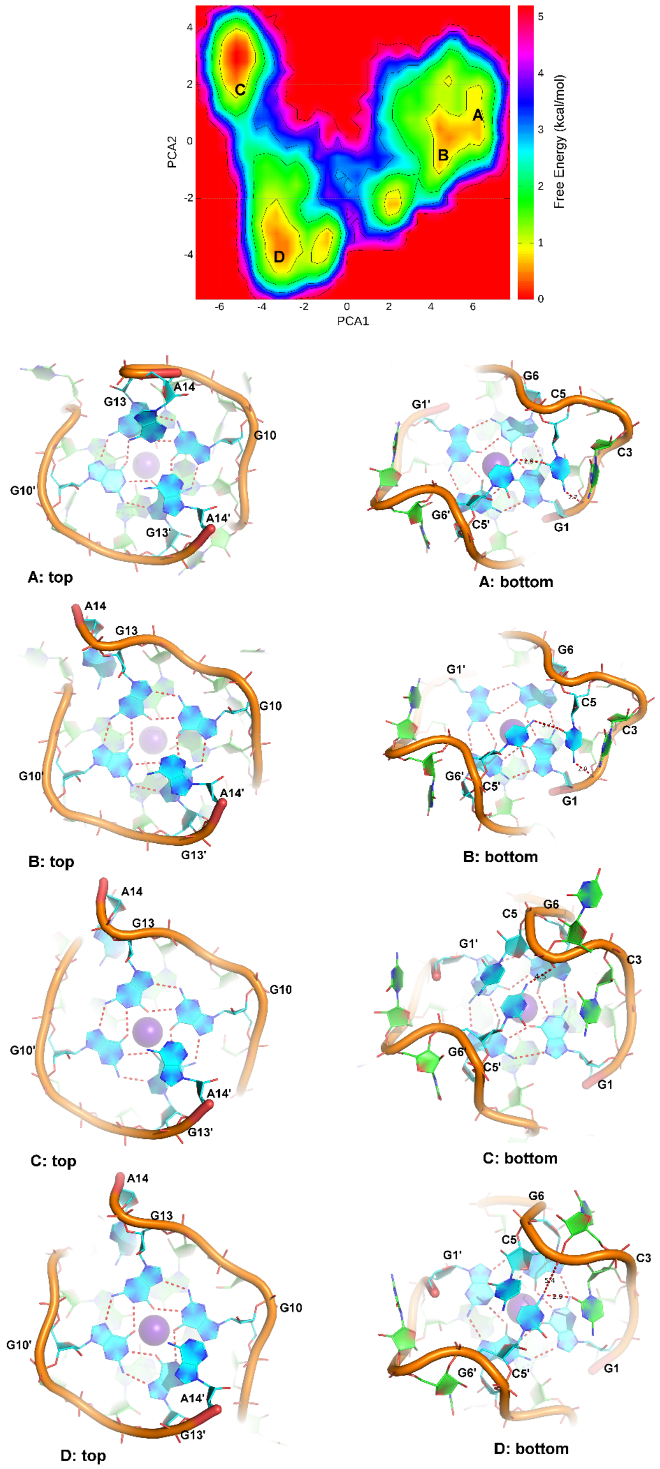


**Figure S2: FES showing the backbone fluctuations of native RNA G4 topology mapped with PCA1 and PCA2**. A variation from open to close-like conformation has been sampled in the long 800 ns of MD simulation showing the flexibility of RNA G4 over a short period of time.

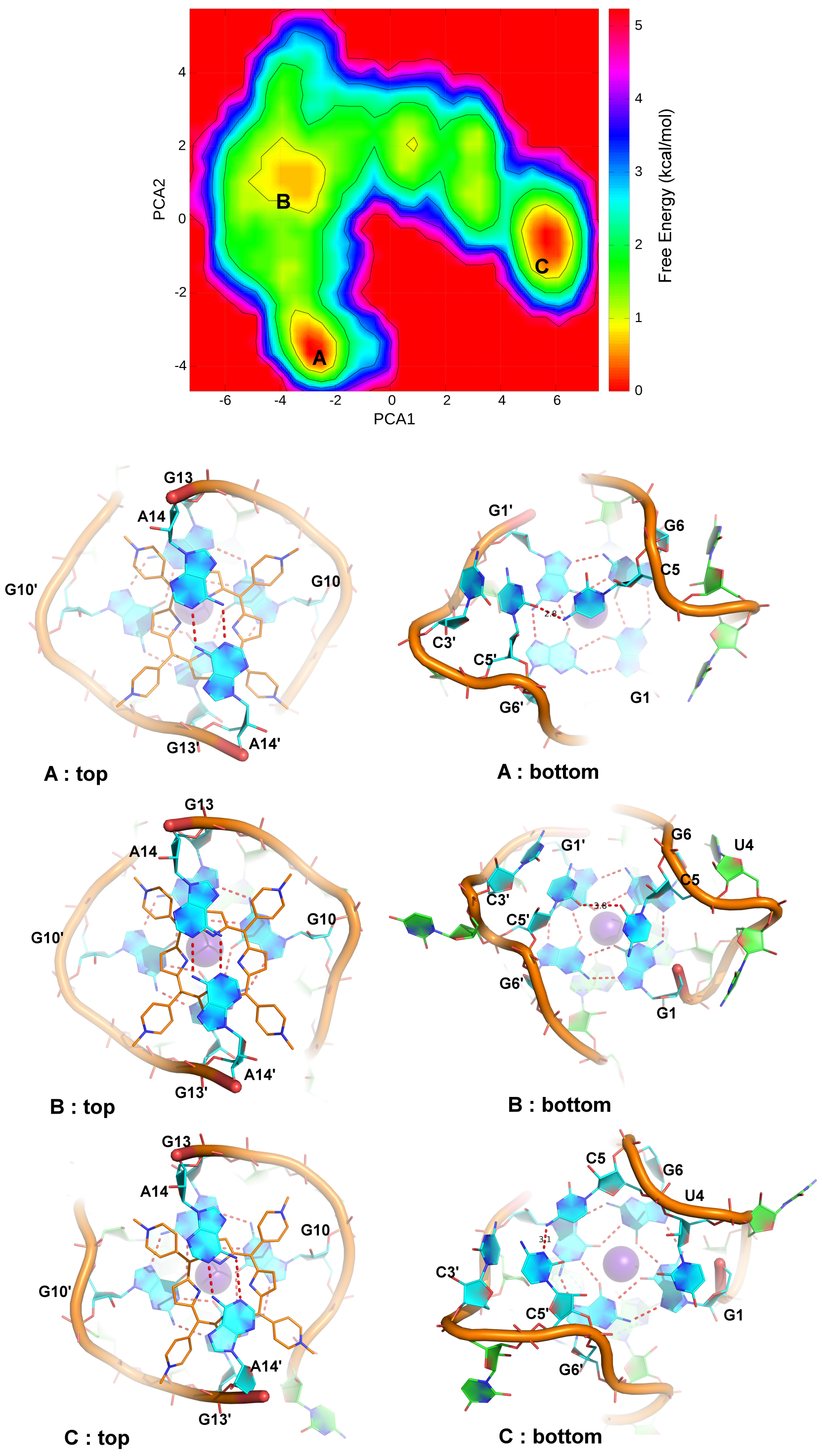


**Figure S3: FES showing the backbone fluctuations of top-face TMPyP4-RNA complex mapped with PCA1 and PCA2.** TMPyP4 mostly seen to stabilize the RNA G4 binding on top-face through stacking interaction with the G-quartet.

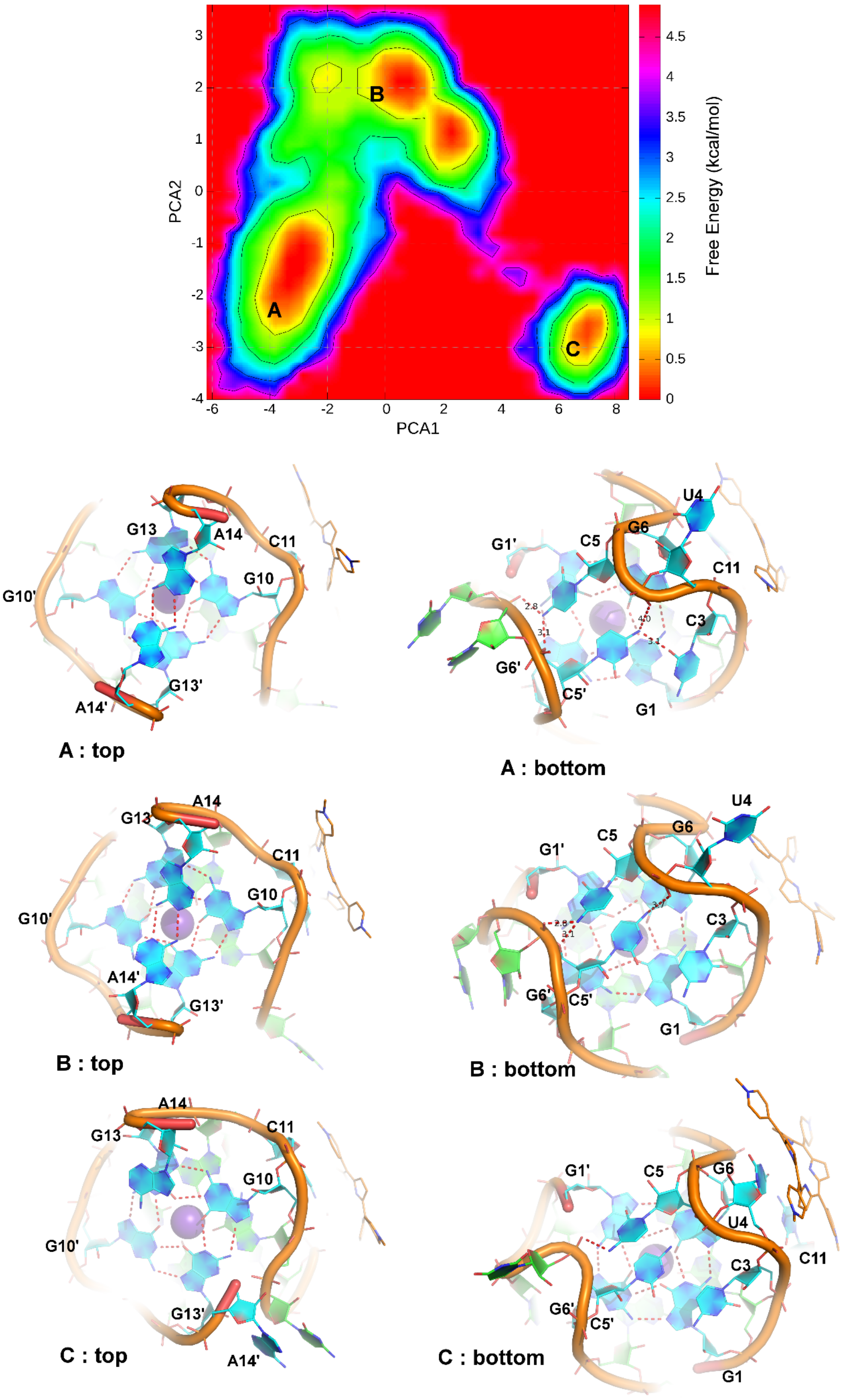


**Figure S4: FES showing the backbone fluctuations of groove bound TMPyP4-RNA complex mapped with PCA1 and PCA2.** TMPyP4 seen to stabilize the RNA G4 through groove binding as RNA G4 topology remains to be in the close-like conformation throughout the 800ns MD simulation.

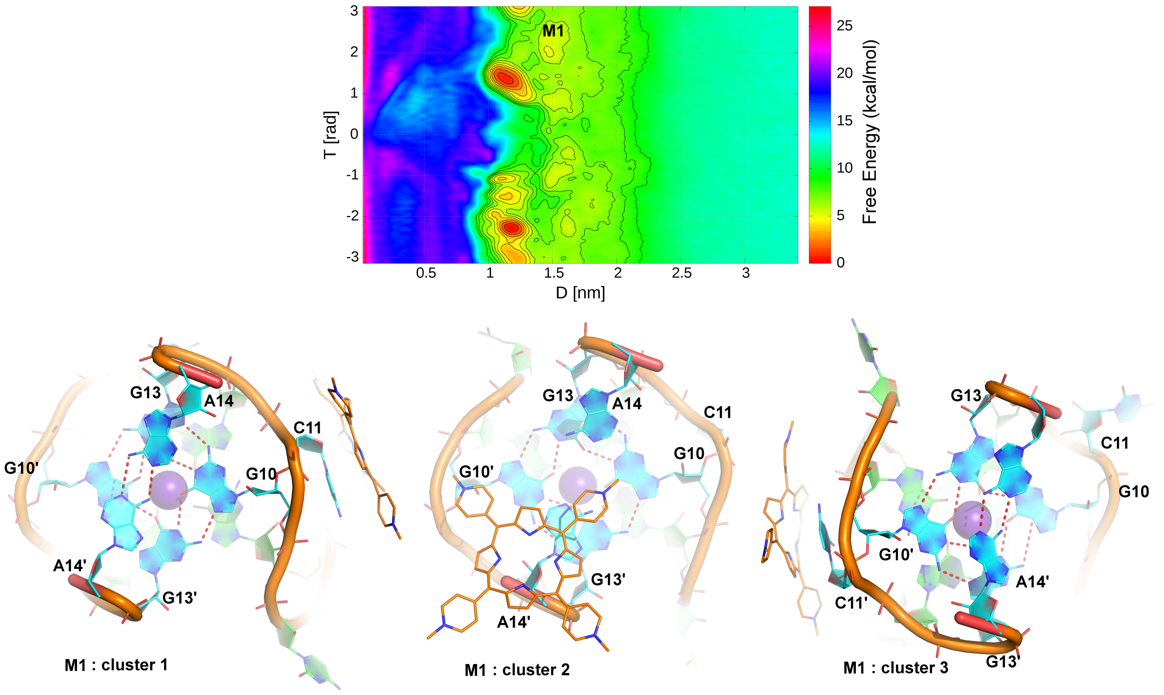


**Figure S5: The FES (D vs T) showing the three most populated clusters of the RNA-TMPyP4 complex on the groove binding position.**

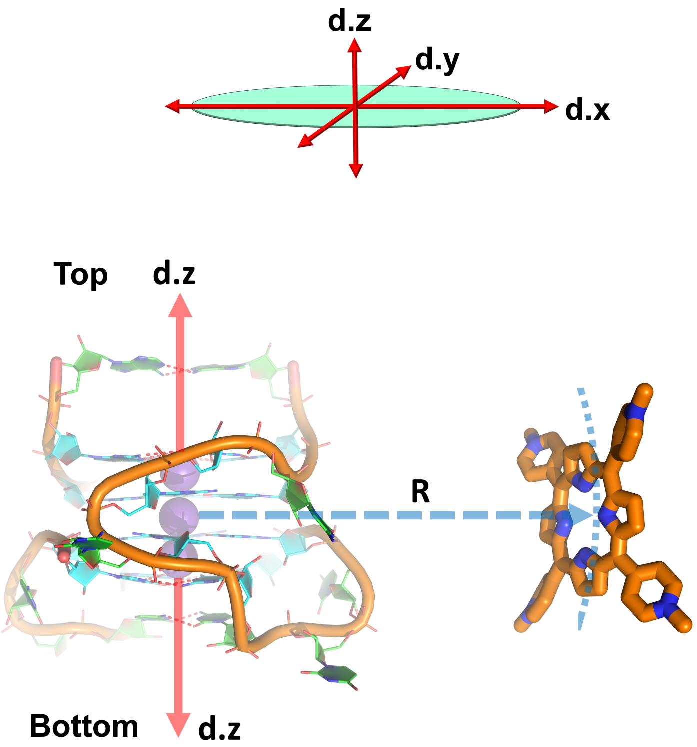


$$R = \sqrt{(d.x)^2 + (d.y)^2}$$

**Figure S6: Construction of the CVs such as POA (Position on the axis) and DFA (Distance from the axis).** POA defines as d.z in nm where the z-axis is aligned with the RNA helix. DFA defines as R in nm where it is calculated as the distance of TMPyP4 from the d.z axis (on the xy plane) thereby describing both the top-face and groove bound states.

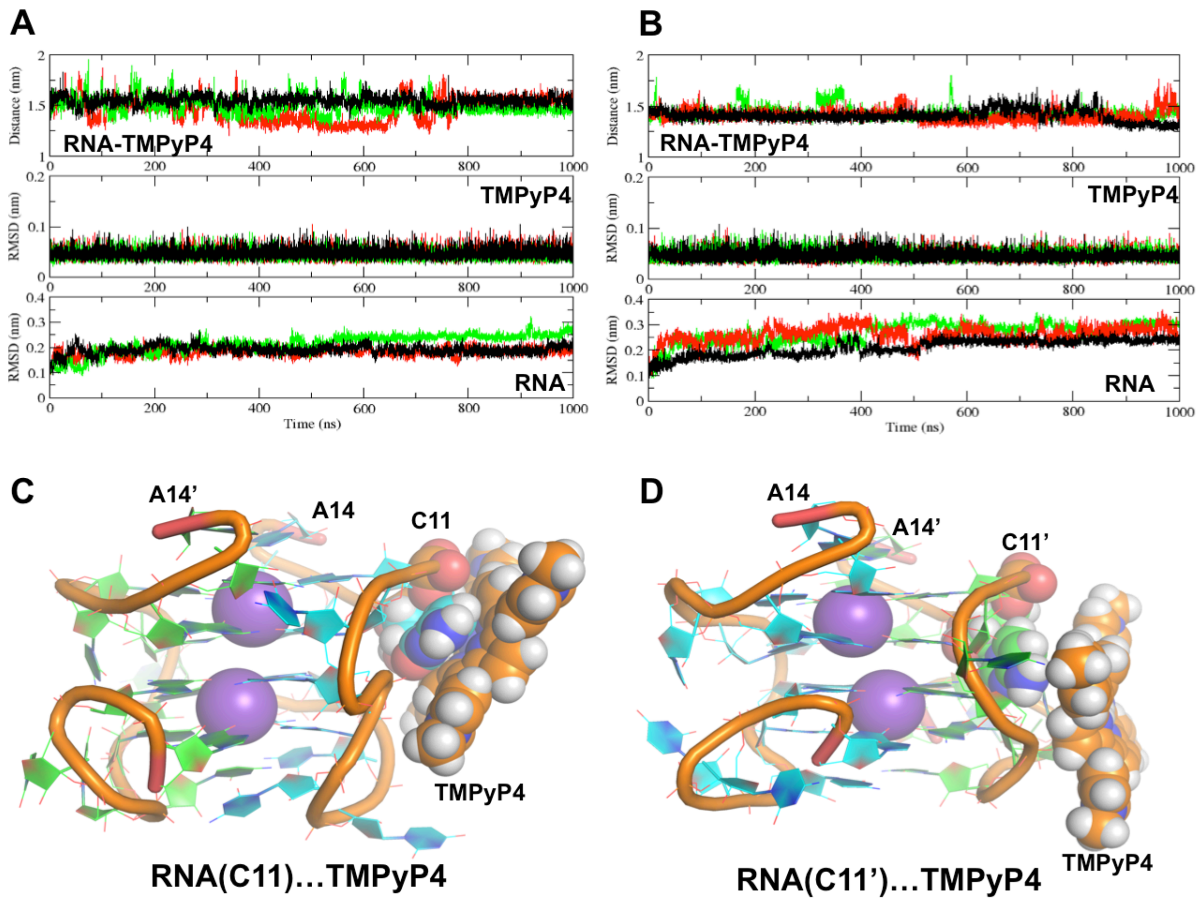


**Figure S7: The MD simulation statistics of TMPyP4-groove bound states.** (a) TMPyP4-RNA(C11) G4: The distance between TMPyP4 and RNA G4 with an average distance of 1.65 nm (top), TMPyP4 RMSD relative to the starting position with an average RMSD being 0.05 nm (middle), and backbone RMSD of RNA G4 relative to the starting position with an average RMSD being 0.19 nm (bottom). (b) TMPyP4-RNA (C11') G4: The distance between TMPyP4 and RNA G4 with an average distance of 1.72 nm (top), TMPyP4 RMSD relative to the starting position with an average RMSD being 0.05 nm (middle), and backbone RMSD of RNA G4 relative to the starting position with an average RMSD being 0.22 nm (bottom). (c) and (d) a representative structure of TMPyP4…RNA G4 complex bound to C11 and C11' groove bound site.

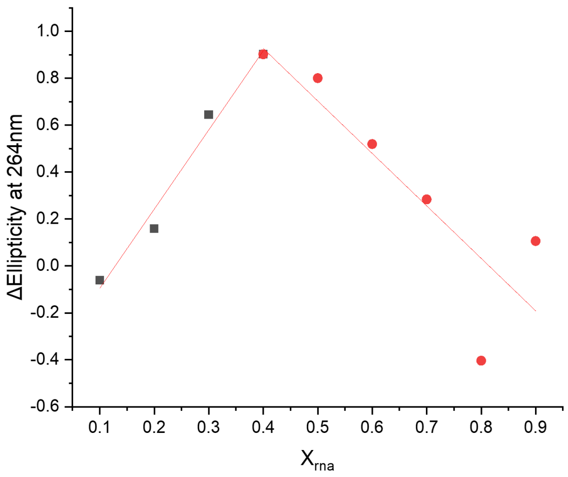


**Figure S8. Example Job plot of RNA PQS18-1 and TMPyP4.** Experiments performed using CD in 10 mM lithium cacodylate buffer, pH 7.0 supplemented with 100 mM of KCl.

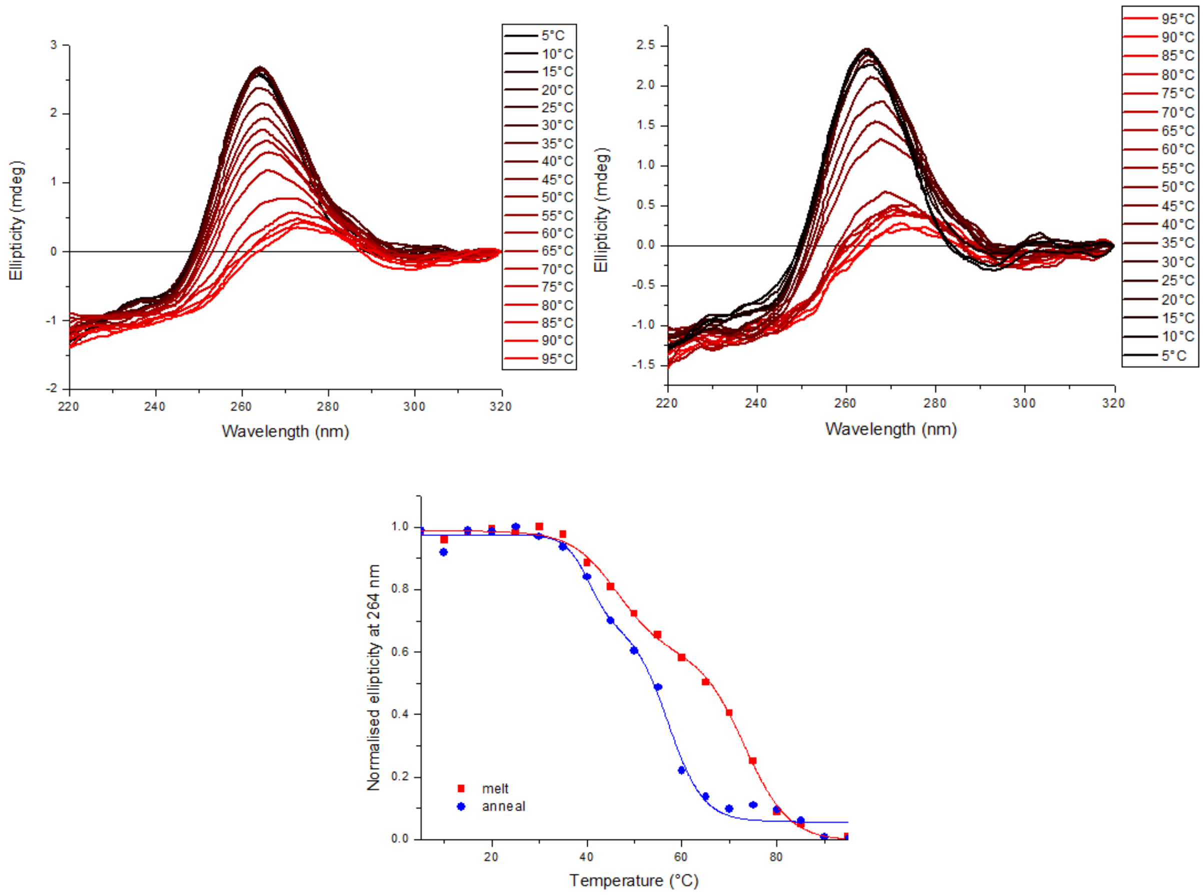


**Figure S9. Example CD melting (Left) and annealing (right) of RNA PQS18-1 and melting/annealing curves (bottom).** Experiments performed at 10 µM RNA in 10 mM lithium cacodylate buffer, pH 7.0 supplemented with 100 mM of KCl.

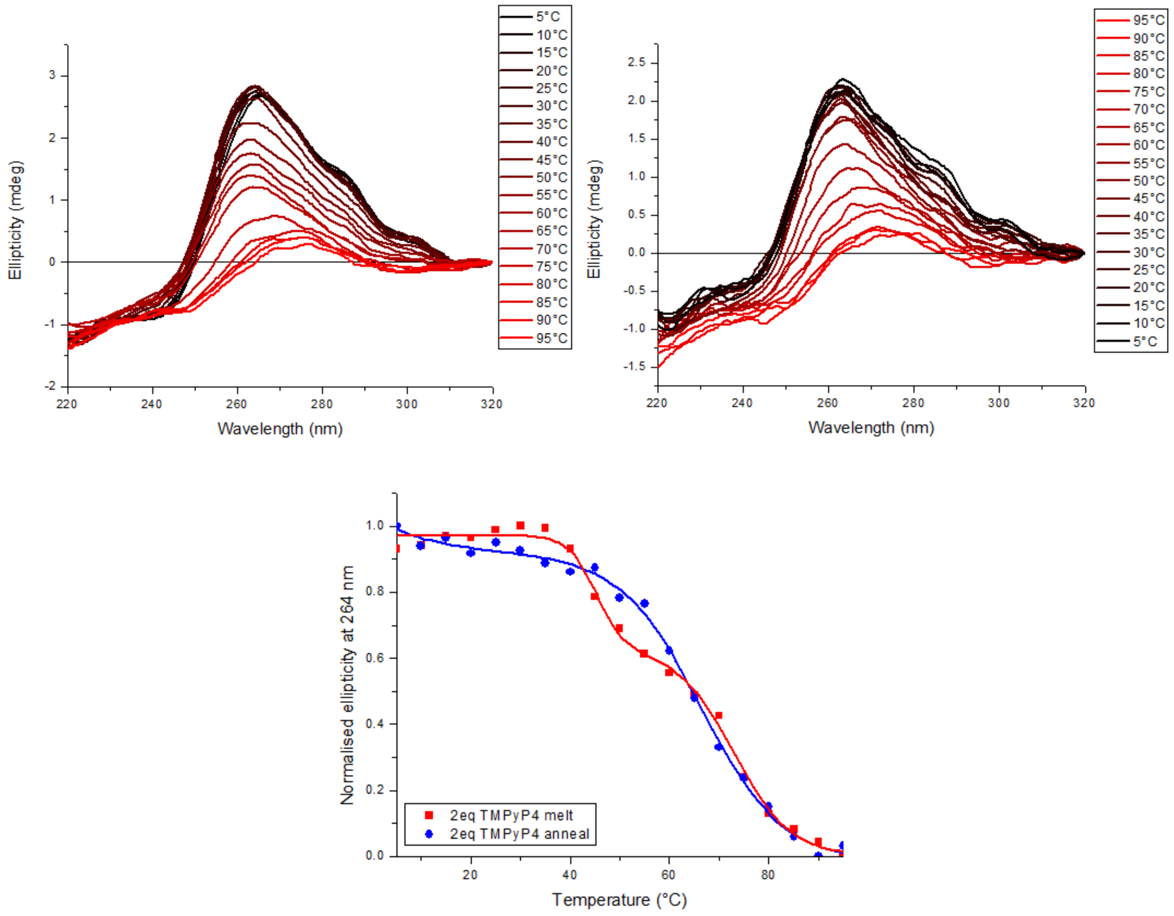


**Figure S10. Example CD melting (Left) and annealing (right) of RNA PQS18-1 and 2 eq TMPyP4 and melting/annealing curves (bottom).** Experiments performed at 10 µM RNA with 20 µM TMPyP4 in 10 mM lithium cacodylate buffer, pH 7.0 supplemented with 100 mM of KCl.

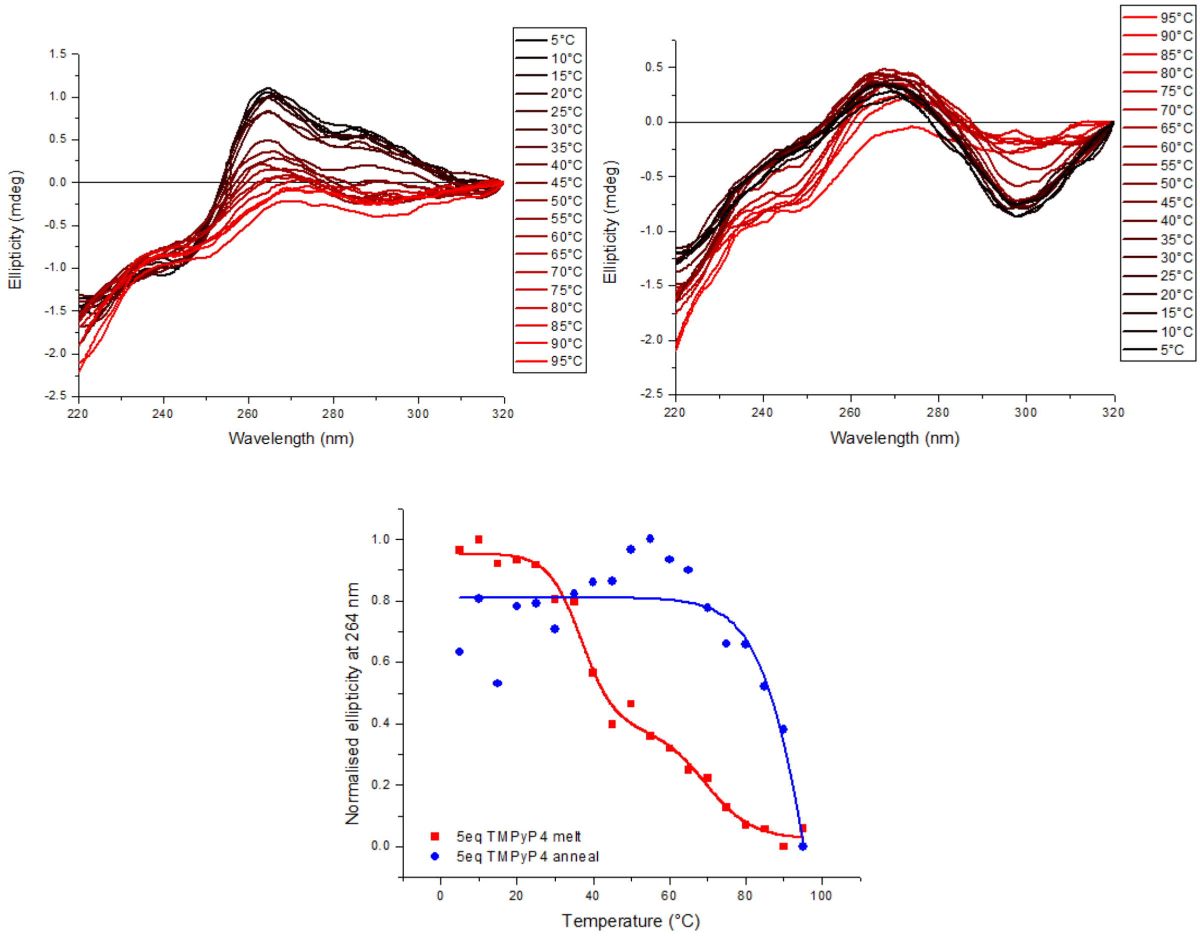


**Figure S11. Example CD melting (Left) and annealing (right) of RNA PQS18-1 and 5 eq TMPyP4 and melting/annealing curves (bottom).** Experiments performed at 10 µM RNA with 50 µM TMPyP4 in 10 mM lithium cacodylate buffer, pH 7.0 supplemented with 100 mM of KCl.

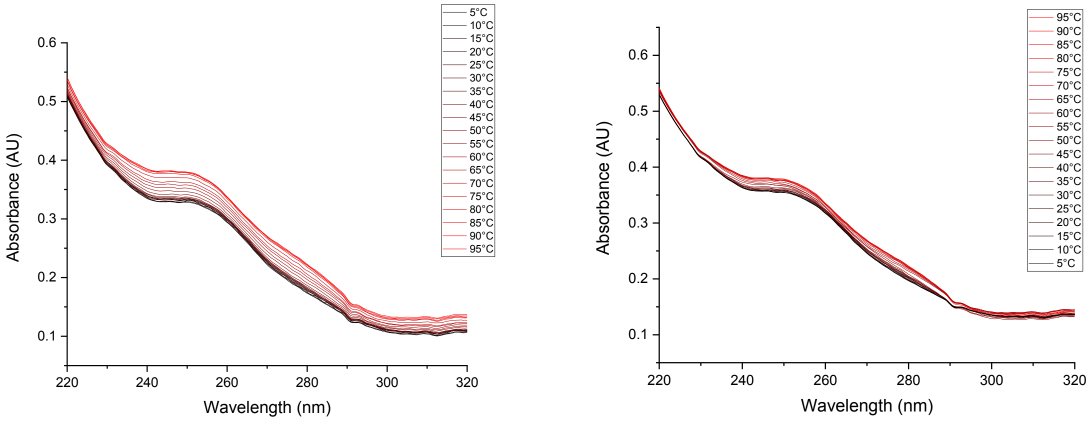


**Figure S12. Example UV melting and annealing of RNA PQS18-1.** Experiments performed at 10 µM RNA in 10 mM lithium cacodylate buffer, pH 7.0 supplemented with 100 mM of KCl.

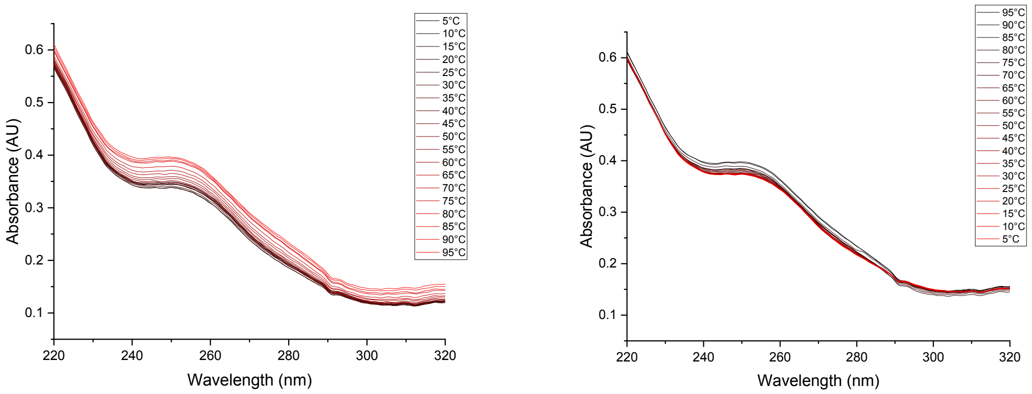


**Figure S13. Example UV melting and annealing of RNA PQS18-1 in the presence of TMPyP4.** Experiments performed at 10 µM RNA in 10 mM lithium cacodylate buffer, pH 7.0 supplemented with 100 mM of KCl and and 2 eq of TMPyP4 (20 µM).

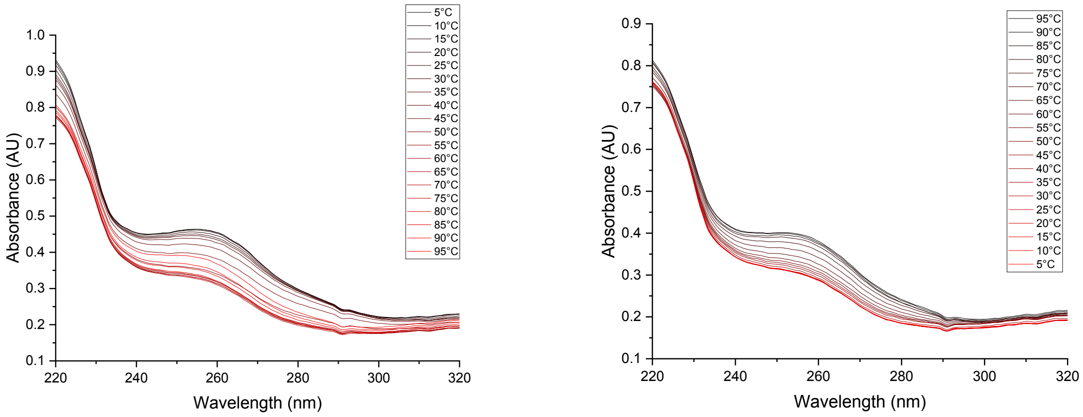


**Figure S14. Example UV melting and annealing of RNA PQS18-1 in the presence of TMPyP4.** Experiments performed at 10 µM RNA in 10 mM lithium cacodylate buffer, pH 7.0 supplemented with 100 mM of KCl and and 5 eq of TMPyP4 (50 µM).

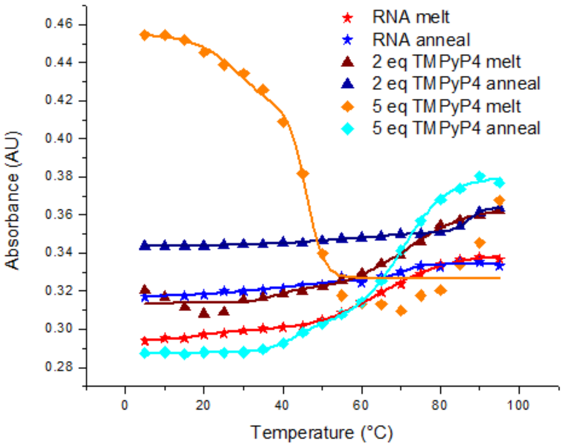


**Figure S15. Example UV melting and annealing of RNA PQS18-1 in the absence and presence of TMPyP4.** Experiments performed at 10 µM RNA in 10 mM lithium cacodylate buffer, pH 7.0 supplemented with 100 mM of KCl and 2 or 5 eq of TMPyP4 (20 and 50 µM respectively) absorbance measured at 260 nm.

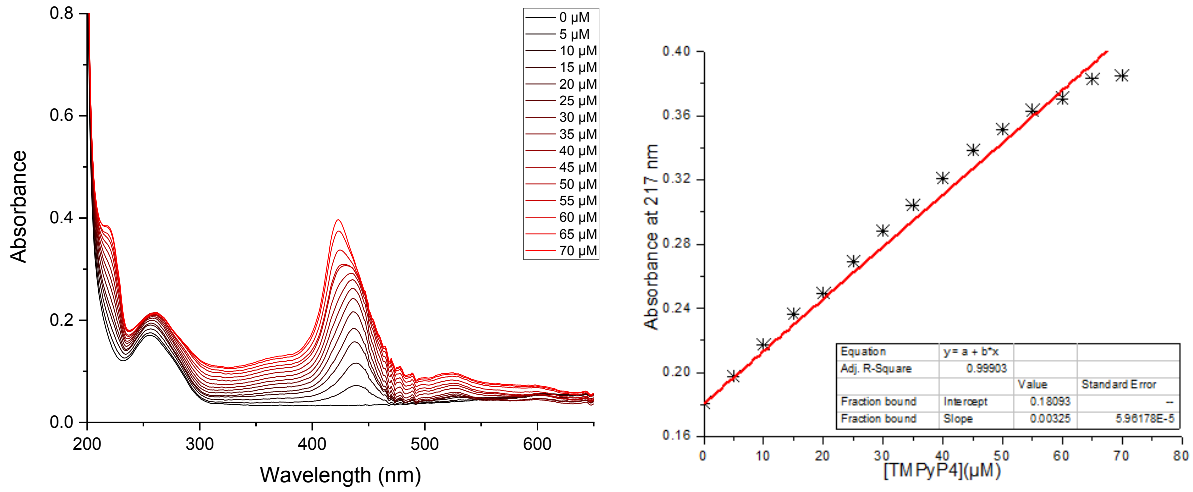


**Figure S16. Example UV titration spectra and plot of UV absorbance at 217 nm of RNA PQS18-1 in the presence of TMPyP4.** Experiments performed at 10 µM RNA in 10 mM lithium cacodylate buffer, pH 7.0 supplemented with 100 mM of KCl and 0 to 70 µM of TMPyP4. Data fitted to a linear function.

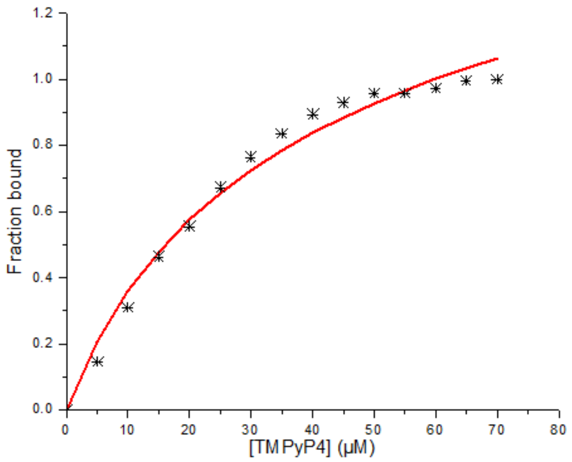


**Figure S17. Example plot of UV absorbance at 440 nm of RNA PQS18-1 in the presence of TMPyP4.** Experiments performed at 10 µM RNA in 10 mM lithium cacodylate buffer, pH 7.0 supplemented with 100 mM of KCl and 0 to 70 µM of TMPyP4. Data fitted to a two inequivalent sites binding model.

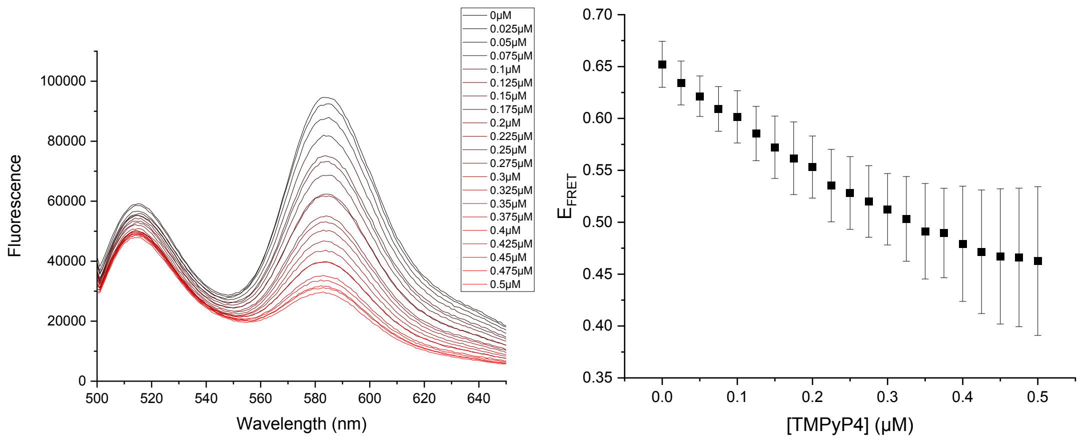


**Figure S18. Example fluorescence spectra of dual-labelled RNA PQS18-1 in the presence of TMPyP4 (left) Relative FRET efficiency with increasing concentration of TMPyP4 (right).** $E_{FRET}$ Experiments performed at 0.1 µM RNA in 10 mM lithium cacodylate buffer, pH 7.0 supplemented with 100 mM of KCl and 0 to 0.5 µM of TMPyP4.

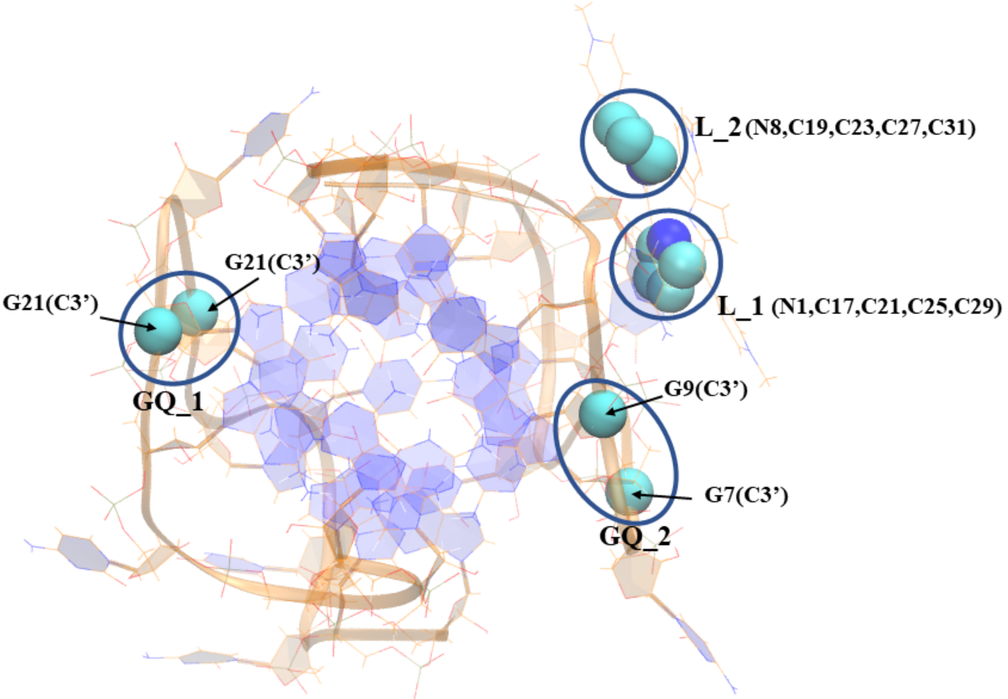


**Figure S19:** The Torsion definition of GQ_1, GQ_2, L_1, L_2 and their corresponding atom definition for the COM representation.

**Table S1: List of atoms with parameters that are involved in building the Distance (D) and Torsion (T) CVs in the WT-MetaD simulation.** The plumed.dat and the input.gro file associated with CVs is also attached in the Supplementary Information. COM stands for Center of Mass. GQ_1, GQ_2 and L_1, L_2 represents two points each from the RNA G4 and TMPyP4, respectively.

| Collective Variables (CV) | Parameters | CVs |
|---|---|---|
| Distance (D) in the WT-MetaD | **RNA G4**: COM{[G7(C3')], [G9(C3')],[G21(C3')],[G23(C3')]}<br>**PMPyP4**: COM (all heavy atoms) | **D**: RNA G4……TMPyP4 |
| Torsion (T) in the WT-MetaD | **GQ_1**: COM{[G21(C3')],[G23(C3')]}<br>**GQ_2**: COM{[G7(C3')], [G9(C3')]}<br>**L_1**: COM{N1,C17,C21,C25,C29}<br>**L_2**: COM{N8,C19,C23,C27,C31} | **T**: GQ_1,GQ_2,L_1,L_2 |

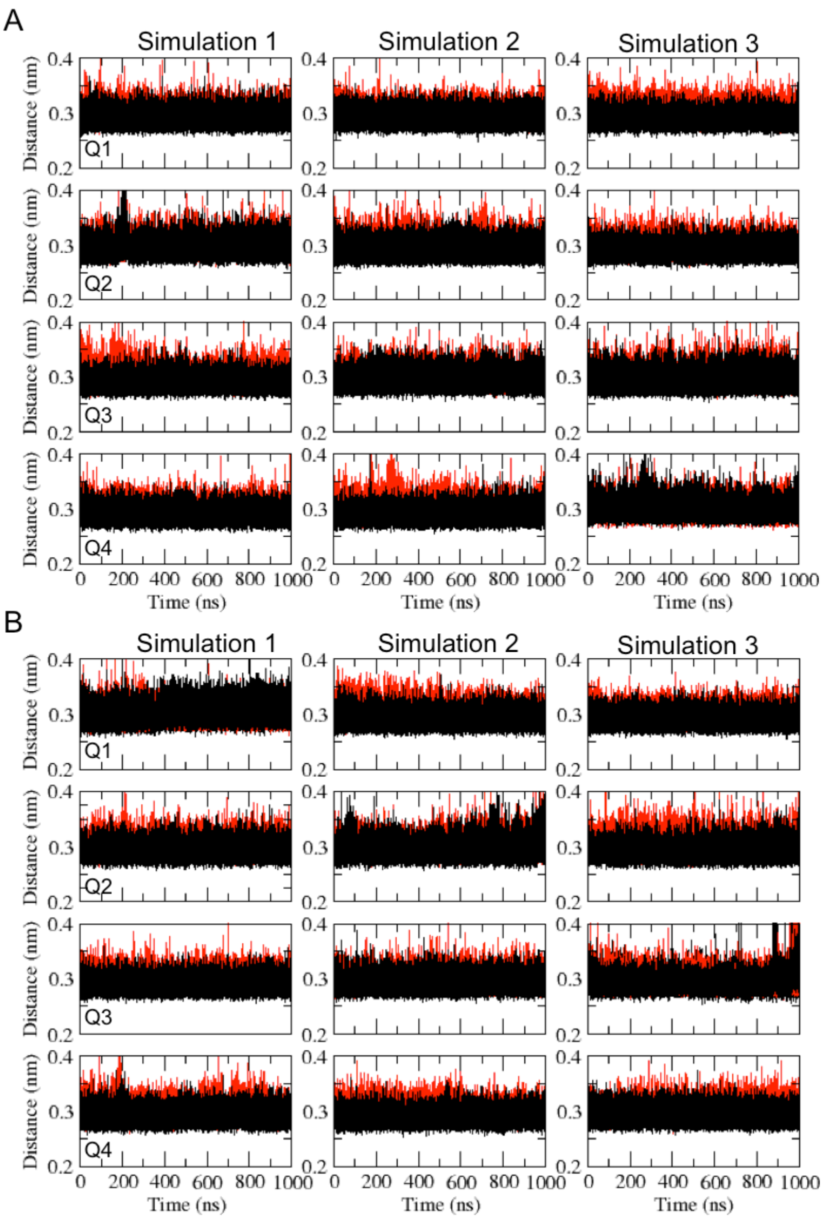


**Figure S20:** Hydrogen bond distances between N1-O6 (black) and N2-N7 (red) atoms in the G4 quartets. The distances have been calculated from the unbiased simulations when TMPyP4 is stacked with (A) C11 and (B) C11' bases. Each row represents a quartet, while each column is a replicate simulation.

**Movie S1:** The dynamics and the rebinding mechanism of TMPyP4 to RNA G4. TMPyP4 is being ctached by C11 base from the bulk and at the end reaching to the top-face bound state.

**Movie S2:** The RNA G4 unfolding mechanism by TMPyP4 through major π-π stacking with Guanine bases from G-tetrads.
